# Molecular Insights into Species-Specific ACE2 Recognition of Coronavirus HKU5

**DOI:** 10.1101/2025.01.10.632062

**Authors:** Yuanyuan Zhang, Yuya Li, Weijie Gao, Dexin Li, Lingyun Xia, Qiang Zhou

## Abstract

Coronaviruses represent a significant zoonotic threat, with host adaptation serving as a pivotal determinant of cross-species transmission. The bat-derived β-coronavirus HKU5 utilizes its spike (S) protein for receptor recognition and viral entry. Here, we report the cryo-electron microscopy (cryo-EM) structure of the HKU5 S protein in a closed conformation. Two fatty acids are found in each protomer of the HKU5 S protein, which stabilize the S protein in the closed conformation. Furthermore, we solve the structure of the HKU5 receptor-binding domain (RBD) in complex with the peptidase domain (PD) of *Pipistrellus abramus* angiotensin-converting enzyme 2 (ACE2), uncovering a unique binding mode distinct from other coronaviruses that use ACE2 as their receptor. Evolutionary and functional analyses indicate that mutations in the RBD can modulate receptor binding, while conservation and structural modeling suggest that HKU5 has the potential to cross the species barrier. Notably, we identify ACE2 orthologs in avian species, such as *Pitta sordida*, that support stable HKU5 RBD binding and interaction. Our functional assays, including pseudovirus entry and cell–cell fusion experiments, demonstrate that HKU5 can exploit ACE2 orthologs across species, providing molecular insights into its host adaptation and underscoring the importance of surveillance for this virus and its zoonotic risk.

## Introduction

Coronaviruses (CoVs) are zoonotic pathogens with significant potential for cross-species transmission, exemplified by SARS-CoV^1^, MERS-CoV^2^ and SARS-CoV-2^3,4^. They often originate from animal reservoirs, especially bats, before spilling over to humans^5–7^. The spike (S) protein of coronaviruses mediates receptor recognition and membrane fusion, with the receptor-binding domain (RBD) typically determining host specificity and spillover potential^8,9^. The RBD can adopt an "up" or "down" conformation, where only the "up" state engages receptors^10–12^. These conformational shifts are regulated by various factors, including sialic acids and fatty acids^13–17^.

Bat coronavirus HKU5 (BatCoV-HKU5) belongs to the β-coronavirus *Merbecovirus* subgenus and is closely related to MERS-CoV^18–20^. Despite being discovered in 2006 and the identification of numerous related sub-clades^21^, HKU5 remains relatively underexplored compared to SARS-CoV, MERS-CoV, and SARS-CoV-2. Recent studies have shown that HKU5 can utilize the angiotensin-converting enzyme 2 (ACE2) from *Pipistrellus abramus* (Japanese house bat), *Homo sapiens* (human) and *Neogale vison* (mink) for host cell entry^22–25^, which serves as a receptor for multiple coronaviruses, including SARS-CoV^26^, SARS-CoV-2^4,27^, HCoV-NL63^28^, MOW15-22^29,30^ and HKU25^31^ and has been implicated in repeated spillover events^32^.

In this work, we determine the cryo-electron microscopy (cryo-EM) structure of the HKU5 S protein in the closed conformation, with each protomer bound with two fatty acids that stabilize this state. We also characterize the interaction between HKU5 RBD and bat ACE2, noting differences from MERS-CoV and SARS-CoV-2. By integrating structural analysis, evolutionary insights, and functional assessment of RBD mutations, we provide a comprehensive perspective on HKU5’s host adaptability and spillover risk. These findings contribute valuable knowledge to coronavirus research, with functional validation of receptor usage demonstrating that HKU5 can engage ACE2 from diverse hosts and underscoring the need for ongoing surveillance of this virus and its zoonotic risk.

## Results

### Structure of HKU5 S protein in the closed conformation

We have over-expressed and purified the HKU5 S protein (UniProt ID: A3EXD0), and then solved its cryo-EM structure at a resolution of 2.4 Å (Supplementary Figs. 1, 2a-d). Unlike the open conformations observed in the S proteins of SARS-CoV-2^11^, MERS-CoV^33^ or HKU1-B^34^, all three RBDs of the HKU5 S protein are in the "down" position (Fig. 1a). This closed conformation hinders interactions with host receptors, potentially delaying host cell entry or facilitating immune evasion^14,35^. Notably, the N174 glycan on the N-terminal domain (NTD) of one protomer forms polar contacts with residues S534, R538, and D549 on the neighboring RBD, which may further stabilize the down state (Supplementary Fig. 2e). In addition, the receptor-binding motif (RBM) is largely buried within the trimer, and the limited exposed surface is masked by glycans from adjacent protomers (N418, N495, and N590), together forming a dynamic glycan shield that reduces the accessibility of the binding interface, a mechanism widely described in coronaviruses.

**Fig. 1.**
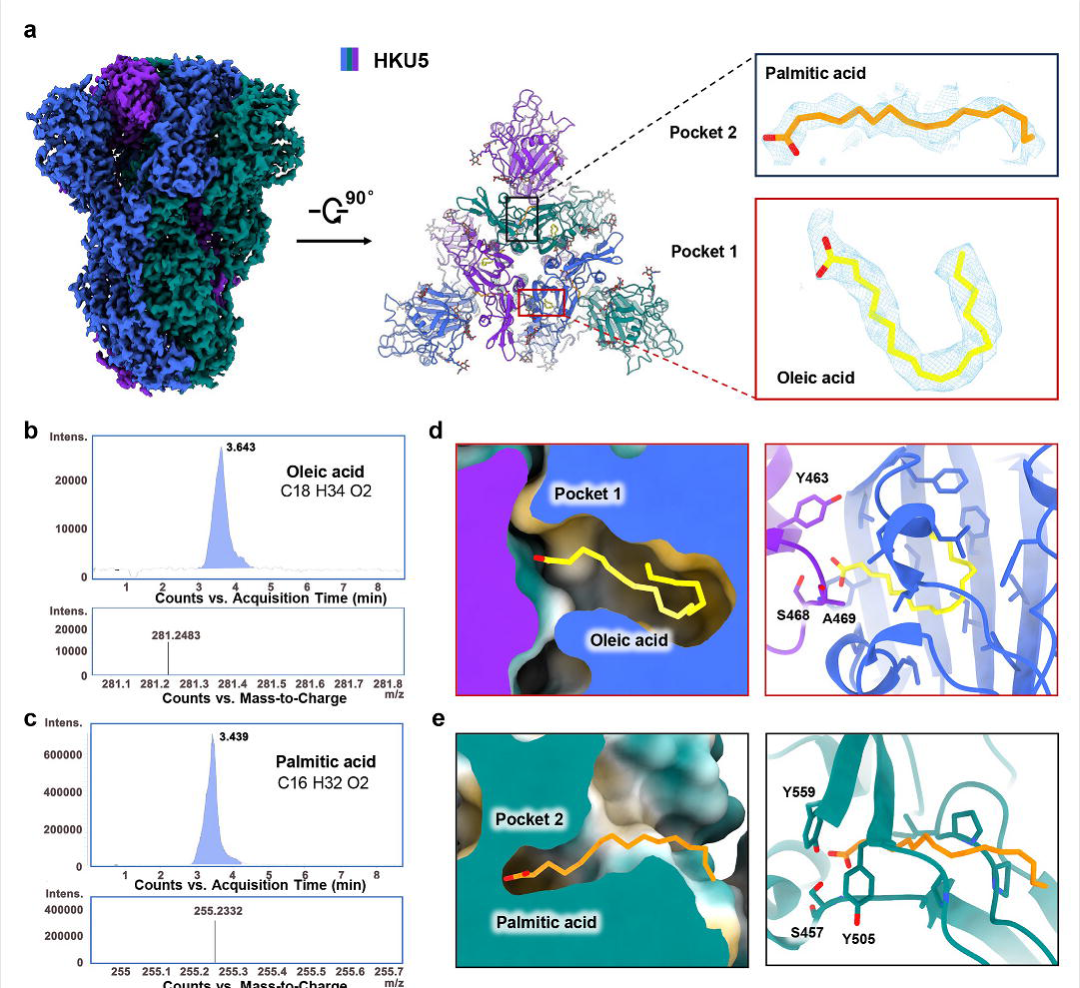
Cryo-EM structure of the HKU5 S protein. **a,** Cryo-EM map shown in side view (Left) at a contour level of 4 σ and the atomic model shown in top view (Middle) of the HKU5 S protein, with protomers displayed in marine blue, purple, and forest green, respectively. Enlarged views of the boxed regions in the middle panel, showing the bound fatty acids (Right), with cryo-EM map shown at a contour level of 3 σ for palmitic acid (orange) and 4 σ for oleic acid (yellow), respectively. **b-c,** Liquid chromatography-mass spectrometry (LC-MS) analysis of purified HKU5 S protein. Top: Chromatograms of C18 and C16 fatty acids. Bottom: Corresponding electrospray ionization time-of-flight (ESI-TOF) mass spectra, with molecular weights of C18H32O2 and C16H34O2 labeled. Intens., intensity; m/z, mass/charge ratio. **d-e,** Binding pockets of oleic acid (**d**) and palmitic acid (**e**) within the HKU5 S protein, highlighting key residues involved in polar interactions, with hydrophobic residues in pocket 1 (F393, L419, L422, L423, F426, V428, F431, P438, L441, L449, and V451, V488, A490, F569, V571) and pocket 2 (P402, P403, I404, A455, and P502). All residues are shown as sticks.

**Fig. 2.**
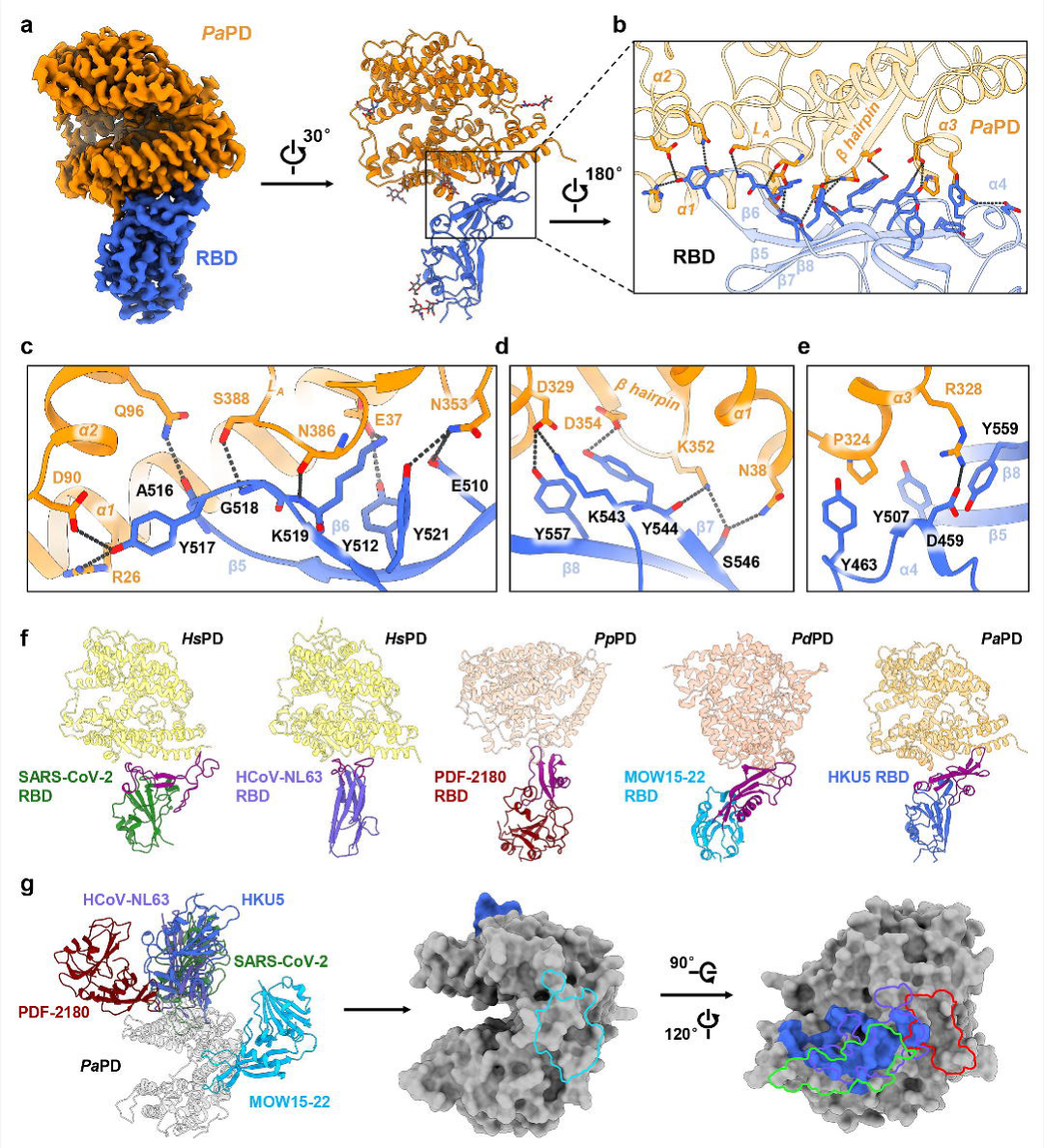
Interactions between HKU5 RBD and bat ACE2. **a,** Cryo-EM map (Left) at a contour level of 4 σ and atomic model shown in cartoon (Right) of HKU5 RBD (marine blue) in complex with *Pa*PD (orange). **b,** Enlarged view of the interaction interface, showing *Pa*PD α1, α2, α3 helices, β-hairpin, and L_A_ loop interacting with β-strands 5∼8 and the α4 of HKU5 RBD. **c-e,** Detailed analysis of the binding interface between HKU5 RBD and *Pa*PD, with polar interactions indicated by dashed black lines. **f,** ACE2 binding modes for SARS-CoV-2, HCoV-NL63, PDF-2180, MOW15-22 and HKU5. *Hs*PD, *Pp*PD, *Pd*PD and *Pa*PD are the peptidase domains of *Homo sapiens*, *Pipistrellus pipistrellus*, *Pteronotus davyi* and *Pipistrellus abramus*, respectively. **g,** Superimposed ACE2 binding sites of SARS-CoV-2, HCoV-NL63, PDF-2180, MOW15-22 and HKU5 on *Pa*PD. For clarity, other PDs are not shown. Binding footprints are shown in cyan (MOW15-22), green (SARS-CoV-2), violet (HCoV-NL63), red (PDF-2180) and marine blue (HKU5), respectively.

In the cryo-EM map of the HKU5 S protein, two additional non-protein densities are identified for each protomer. The first one is in a pocket (pocket 1) near the interface between the RBDs of adjacent protomers, while the second one is in another pocket within the RBD (pocket 2), near the RBM (Fig. 1a and Supplementary Fig. 2e). Mass spectrometry experiments suggest the HKU5 S protein contains oleic acid and palmitic acid (Fig. 1b, c and Supplementary Fig. 3), which are fitted to pocket 1 and 2 based on their density shape, respectively (Fig. 1a). The pocket 1 is formed by hydrophobic residues F393, L419, L422, L423, F426, V428, F431, P438, L441, L449, V451, V488, A490, F569, and V571 on the RBD (Fig. 1d). The polar head of oleic acid interacts with residues Y463, S468, and A469 from an adjacent protomer (Fig. 1d). In a cryo-EM map of the HKU5 S protein with S468A&Y463A double mutant (S_DM1_ protein), the density corresponding to oleic acid was absent, indicating the importance of these polar residues for the interaction with oleic acid (Supplementary Figs. 4, 5). Compared to the wild-type (WT) protein, the S_DM1_ protein exhibited a ∼5° clockwise rotation of the NTD (Supplementary Fig. 5a and Supplementary Movie 1), resulting in a looser trimeric assembly^36^. Additionally, the SD1 and SD2 domains shifted away from the RBD toward the NTD, forming new polar interactions such as a salt bridge between R689 of SD2 and D346 of NTD, and a hydrogen bond network involving residues ^689^RMT^691^ of SD2 and ^348^GYDD^351^ of NTD (Supplementary Fig. 5b-e and Supplementary Movie 2). These changes were accompanied by structural remodeling, suggesting that oleic acid is beneficial for maintaining the closed conformation of the S protein. This observation parallels findings in SARS-CoV-2, where linoleic acid binding stabilizes the closed state of the S protein^17^. The pocket 2, where the palmitic acid binds, is in the interior of the RBD (Fig. 1e), which has not been reported in any other S protein of coronaviruses. Although the pocket 2 is close to the receptor-binding interface, it does not directly contact it, possibly suggesting a structural or regulatory role in modulating RBD dynamics rather than directly influencing ACE2 binding. To further examine functional effects, we generated pseudoviruses carrying either WT S or the fatty acid binding site mutants, S_DM1_ and S_DM2_ (S457A&Y559A). Infection assays revealed no significant differences in entry efficiency (Supplementary Fig. 5f, g). Together, these results suggest that fatty acid binding plays a role in stabilizing the closed conformation of the S protein, but is not essential for ACE2-mediated entry.

### Interaction of HKU5 RBD with bat ACE2

To elucidate the interaction between the HKU5 S protein and ACE2 of its natural host, *Pipistrellus abramus*, we conducted biochemical and structural studies. The S protein in its closed conformation does not co-migrate with the peptidase domain (PD) of the *Pipistrellus abramus* ACE2 (*Pa*PD) in gel filtration chromatography, indicating that the closed S trimer cannot directly bind to the receptor (Supplementary Fig. 6). To understand the molecular mechanism underlying HKU5 S protein receptor recognition, we determined the cryo-EM structure of the HKU5 RBD in complex with *Pa*PD at a resolution of 3.2 Å, allowing accurate modeling and analysis (Fig. 2a and Supplementary Fig. 7a-d).

The RBM of the HKU5 RBD adopts a compact super-secondary structure comprising a four-stranded anti-parallel β-sheet (β5∼β8, residues N504∼P560) and a short helix (α4, residues S457∼L464), which are critical for receptor recognition and binding (Fig. 2a, b and Supplementary Fig. 7e). The *Pa*PD comprises key structural elements that facilitate its interaction with HKU5 RBD, including the α1 helix (18∼54 a. a.), α2 helix (90∼101 a. a.), α3 helix (322∼330 a. a.), a β-hairpin (346∼358 a. a.), and a loop region (L_A_; 384∼399 a. a.) (Fig. 2a, b).

The interface between the RBD and *Pa*PD spans an area of ∼1107 Å² and can be divided into three patches, each playing a pivotal role in stabilizing the complex. The patch 1 involves interactions between RBD residues from β5 and β6 (E510, Y512, A516, Y517, G518, K519, Y521) and *Pa*PD residues in the α1 helix (R26, E37), α2 helix (D90, Q96), β-hairpin (N353), and L_A_ loop (N386, S388) (Fig. 2c). The interaction is mediated by eight hydrogen bonds and one salt bridge (Fig. 2c), which collectively ensure the stability of this initial contact point. The predominance of polar and charged residues highlights the importance of this region in the high-affinity binding observed in the HKU5 RBD–*Pa*PD complex. The residues of patch 2 from β7 and β8 of the RBD (K543, Y544, S546, Y557) interact with *Pa*PD residues from the α1 helix (N38), α3 helix (D329), and β-hairpin (K352, D354) (Fig. 2d). Notably, K543 and Y557 in RBD form a salt bridge and a hydrogen bond with D329 in *Pa*PD, respectively, while Y544 in RBD interacts with D354 and K352 in *Pa*PD. N38 and K352 in *Pa*PD also engage with S546 in RBD through hydrogen bonds (Fig. 2d). Patch 3 incorporates a variety of stabilizing interactions (Fig. 2e). Specifically, D459 in the RBM and Y559 in β8 of the RBD form a salt bridge and a cation-π interaction with R328 in the *Pa*PD α3 helix, respectively. Additionally, hydrophobic interactions are observed in the region encompassing Y463 of the RBD α4, Y507 of RBD β5, and P324 of the *Pa*PD α3 helix. Collectively, these three patches establish a highly specific binding interface between the HKU5 RBD and *Pa*PD, which is crucial for host ACE2 recognition and initiation of viral entry.

In addition to HKU5, numerous other coronaviruses utilize ACE2 for host cell entry, including the epidemic- or pandemic-causing SARS-CoV^26^ and SARS-CoV-2^4,27^, as well as human coronavirus HCoV-NL63^28^, and several bat coronaviruses, such as NeoCoV^37^, PDF-2180^37^, PRD-0038^38^, and MOW15-22^29,30^. The RBM of SARS-CoV-2 forms an accessible, expansive surface that tightly binds human ACE2. HCoV-NL63 RBD has a β-sandwich core with three non-contiguous RBMs clustered at the center. The RBMs of PDF-2180 and MOW15-22 adopt a four-stranded anti-parallel β-sheet similar to HKU5, but HKU5 is more compact with an extended fourth strand (Fig. 2f and Supplementary Fig. 7e).

Structural comparison reveals differences of the HKU5 RBD–ACE2 interface to the other coronaviruses (Fig. 2g). HKU5 shares partial binding footprints with the known modes of SARS-CoV-2, HCoV-NL63, and PDF-2180, but lacks conserved residues among these viruses (Fig. 2g). This indicates that the HKU5 RBD employs a unique interface to interact with ACE2, likely contributing to the host specificity.

### Evolutionary dynamics of HKU5 RBD

To explore the evolutionary dynamics of the HKU5 RBD, we performed a clustering analysis of the sequences from the NCBI protein database sharing more than 80% identity with the HKU5 S protein (Supplementary Fig. 8). The results show that six HKU5 sub-clades recorded in the NCBI Taxonomy database are distributed across two clusters (Supplementary Fig. 8). Most of the HKU5 RBD residues are highly conserved across various sub-clades, particularly within the core framework of the RBD, indicating their critical role in maintaining structural integrity (Supplementary Figs. 8, 9). However, over 30 variable sites are found in the RBM, particularly spanning the segment Q494∼G564 (Supplementary Figs. 8, 9) that accounts for more than two-thirds of the residue indels and point mutations.

To evaluate the impact of different HKU5 sub-clades on ACE2 recognition specificity, we employed bio-layer interferometry (BLI) to quantify the *Pa*PD binding affinity of various HKU5 RBD variants (Fig. 3a). The highest affinity is observed for HKU5, with a KD of 19.94 ± 0.23 nM. BtPa-BetaCoV/GD2013 (GenBank accession: AIA62343.1) and BatCoV-HKU5-3 (GenBank accession: ABN10893.1) display similar binding affinity, at 23.40 ± 0.27 nM and 25.17 ± 0.29 nM, respectively. BatCoV-HKU5-related (GenBank accession: QHA24687.1) viruses show a slightly lower affinity, with a KD of 35.24 ± 0.37 nM. These HKU5 sub-clades exhibit *Pa*PD binding affinity either comparable to or slightly lower than that of the HKU5, suggesting that the mutations observed in these variants have only a minor effect on receptor binding.

**Fig. 3.**
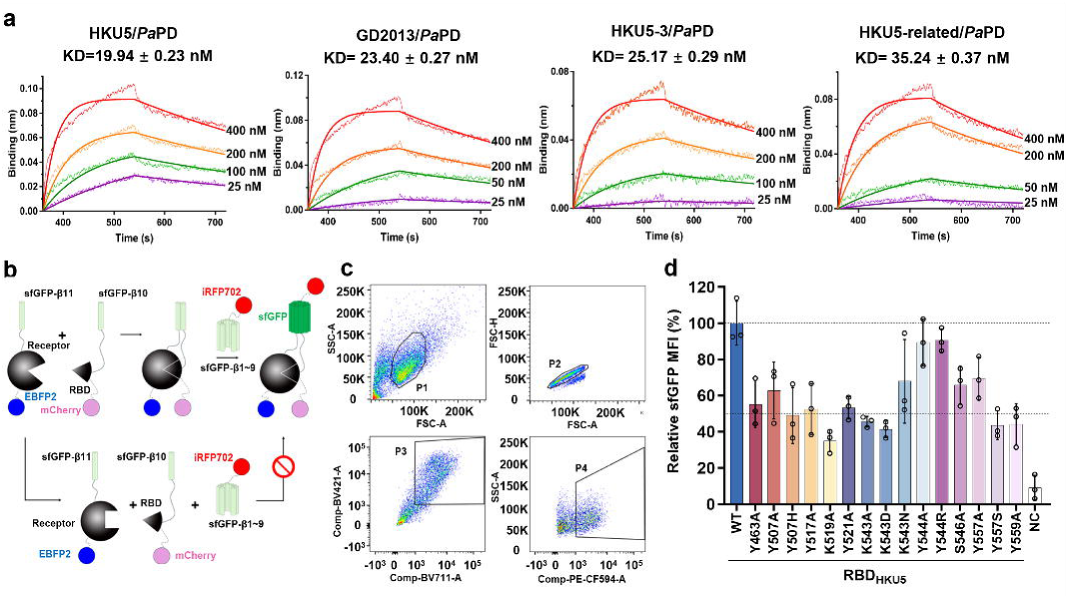
Interaction and functional analysis of HKU5 sub-clades RBDs with *Pa*PD. **a,** Binding and dissociation kinetics of RBDs of four HKU5 sub-clades (BatCoV-HKU5 (UniProt ID: A3EXD0), BatCoV-HKU5-3 (GenBank accession: ABN10893.1), BtPa-BetaCoV/GD2013 (GenBank accession: AIA62343.1), and BatCoV-HKU5-Related (GenBank accession: QHA24687.1)) with *Pa*PD measured by bio-layer interferometry (BLI). R² values are 0.9776, 0.9830, 0.9867, and 0.9861, respectively. **b,** Experimental schematic of tripartite split-fluorescence system to evaluate HKU5 RBD-*Pa*PD interactions. **c,** Flow cytometry-based sorting strategy for single-cell populations co-expressing HKU5 RBD and *Pa*PD. The first gate (P1) was applied to exclude cell debris, followed by the second gate (P2) to remove doublets or adhered cells. Populations P3 and P4 were subsequently defined based on fluorescence intensity, and cells in P3 and P4 were isolated. sfGFP fluorescence intensity was quantified in the P4 population. **d,** Relative sfGFP fluorescence intensity for WT or mutant HKU5 RBDs binding *Pa*PD, normalized to the WT signal. Data represent mean ± standard deviation (SD) from three independent experiments (n = 3). Source data are provided as a Source Data file.

To further investigate the influence of conserved interface residues on the interaction between HKU5 RBD and *Pa*PD, we utilized a tripartite split-fluorescence system^39^ in conjunction with flow cytometry to identify potential key residues within the HKU5 RBD (Fig. 3b, c). These assays showed high sensitivity for the interaction of the HKU5 RBD with *Pa*PD, with signal intensity exceeding that of the HKU5 RBD with PD of *Homo sapiens* ACE2 (*Hs*PD), which is a negative control, by more than tenfold (Fig. 3d). Targeted mutations of interface residues revealed that, except for Y544, mutations of the other residues, namely Y463, Y507, Y517, K519, Y521, K543, S546, Y557, and Y559, led to varying degrees of destabilization of the interaction, as evidenced by decreased fluorescence (Fig. 3d). Notably, the K519A mutation caused a reduction in fluorescence intensity by more than 50%. It is important to highlight that residues Y507, K543, Y544, and Y557 are not highly conserved among HKU5 sub-clades (Fig. 3d). When these non-conserved residues were mutated to their corresponding residues in other sub-clades, we observed a reduction in fluorescence intensity by approximately 50%, as in the cases of Y507H, K543D, and Y557S (Fig. 3d). These findings contribute to a deeper understanding of the factors influencing the binding affinity of the HKU5 virus for its receptor.

### Cross-species transmission risk analysis of HKU5

To investigate the potential for cross-species transmission of HKU5, we conducted a detailed structural analysis of the *Pa*PD residues recognized by HKU5. We introduced point mutations at several polar residues of *Pa*PD that are likely to stabilize the interface, including R26A, E37A, Q96A, N321A, P324A, W327A, R328A, D329A, K352A, N353A, D354A, and S388A (Fig. 4a). Tripartite split-fluorescence experiments performed on these mutants revealed that most of the mutations resulted in a noticeable reduction in relative fluorescence intensity, indicating their significant contribution to the virus–receptor interaction (Fig. 4b). Notably, the R328A mutation in *Pa*PD caused a reduction in relative fluorescence intensity to below 50%. The R328 in *Pa*PD stabilizes the interface through a salt bridge with D459 and a cation-π interaction with Y559 in HKU5 RBD, thus maintaining the interaction between HKU5 and *Pa*PD (Fig. 2e). This result aligns with the reduced fluorescence intensity observed in the HKU5 RBD Y559A mutant (Figs. 3d, 4b). Collectively, the W327A, R328A, D329A, K352A, N353A, and D354A mutations resulted in a marked reduction of both fluorescence intensity and pseudovirus entry efficiency (Fig. 4b, c and Supplementary Fig. 11b). These residues are located within the core of the receptor-binding motif, and the loss of function upon mutation highlights their critical role in mediating the interaction between the HKU5 RBD and ACE2. Sequence alignment of ACE2 orthologs from representative species revealed that, although ACE2 is generally conserved among vertebrates, the key receptor-binding motifs (^327^WRD^329^ and ^352^KND^354^) are more conserved in avian species, exemplified by *Pitta sordida* (Hooded Pitta) (Fig. 4d and Supplementary Fig. 10). The sequence identity of the PD of *Pitta sordida* ACE2 (*Ps*PD) with *Pa*PD and *Hs*PD is 69% and 76%, respectively. Both pseudovirus infection and cell–cell fusion assays demonstrated that Lenti-X 293T or Caco-2 cells stably expressing *Pitta sordida* ACE2 (*Ps*ACE2) could robustly support HKU5 entry (Fig. 4e, f and Supplementary Fig. 11). We measured the binding affinity of *Ps*PD for HKU5 RBD by BLI, which yielded a dissociation constant (KD) of 122.9 ± 1.3 nM (Supplementary Fig. 12).

**Fig. 4.**
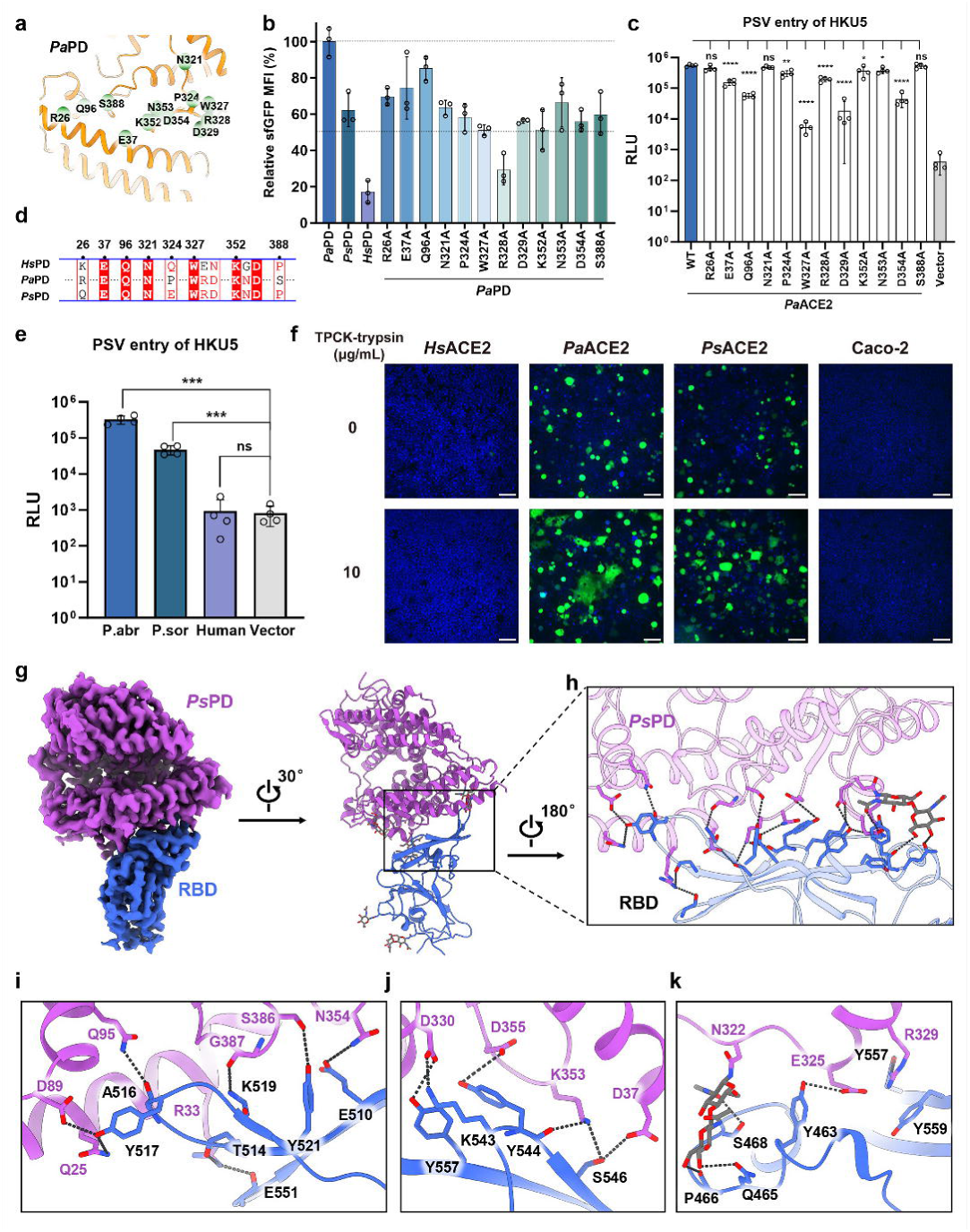
Mutational analysis of ACE2 interface residues and cross-species RBD interactions. **a**, Critical interface residues of *Pa*PD involved in HKU5 RBD binding. **b,** Effects of *Pa*PD, *Ps*PD, *Hs*PD and *Pa*PD mutants on HKU5 RBD binding, evaluated by flow cytometry and tripartite split-fluorescence-based assays. The sfGFP fluorescence intensity was normalized to WT *Pa*PD. Data represent mean ± SD from three independent experiments (n = 3). **c,** HKU5 S pseudovirus (PSV) entry into Lenti-X 293T cells transiently expressing the indicated *Pipistrellus abramus* ACE2 (*Pa*ACE2) mutants. Data represent mean ± SD of four technical replicates from one representative experiment (three independent experiments performed in total). **d,** Conservation analysis of *Pipistrellus abramus* (*Pa*PD) interface residues in *Homo sapiens* (*Hs*PD) and *Pitta sordida* (*Ps*PD), highlighting conserved residues essential for cross-species interactions. **e,** Entry of HKU5 S pseudovirus (PSV) into Lenti-X 293T cells stably transfected with three ACE2 orthologs, *Pipistrellus abramus* (P.abr), *Pitta sordida* (P.sor) and *Homo sapiens* (human). Data represent mean ± SD of four technical replicates from one representative experiment (three independent experiments performed in total). **f,** HKU5 S-mediated cell–cell fusion using Caco-2 cells stably expressing the indicated ACE2 orthologs, *Homo sapiens* (*Hs*ACE2), *Pipistrellus abramus* (*Pa*ACE2) and *Pitta sordida* (*Ps*ACE2), with or without the addition of 10 μg/mL TPCK-treated trypsin. Untransfected Caco-2 cells under the same conditions served as the blank control. Representative images from three independent experiments showing similar results. **g,** Cryo-EM map (Left) at a contour level of 4 σ and atomic model shown in cartoon (Right) of HKU5 RBD (marine blue) in complex with *Ps*PD (purple). **h,** Enlarged view of the interaction interface between HKU5 RBD and *Ps*PD. **i-k,** Detailed analysis of interface interactions between *Ps*PD and HKU5 RBD, with polar interactions indicated by dashed black lines. Statistical significance was determined using two-sided unpaired Student’s *t*-tests. ns, not significant (P ≥ 0.05); * P < 0.05; ** P < 0.01; *** P < 0.001; **** P < 0.0001. Source data are provided as a Source Data file.

To analyze the impact of the receptor interface residue changes on HKU5 RBD recognition, we solved the cryo-EM structure of the HKU5 RBD–*Ps*PD complex. Likely due to the reduced affinity, a large fraction of the *Ps*PD molecules remained unbound in the 2D classification. However, after collecting sufficient data, we solved the HKU5 RBD–*Ps*PD complex at a resolution of 3.2 Å (Fig. 4g and Supplementary Fig. 13). *Ps*PD binds to the HKU5 RBD in a manner similar to that of *Pa*PD (Supplementary Figs. 14, 15). The core interaction interface between *Ps*PD and HKU5 RBD is maintained by the WRD and KND motifs in *Ps*PD, along with other conserved residues (Fig. 4h-k). Some interface residues of *Ps*PD were mutated relative to *Pa*PD, including Q25, D37, E325, G387, and P389, which correspond to R26, N38, P324, N386, and S388 in *Pa*PD, respectively (Fig. 4d and Supplementary Fig. 15). The E325 in *Ps*PD (corresponding to P324 in *Pa*PD) disrupts its hydrophobic interaction with Y463 in RBD but forms a hydrogen bond instead (Fig. 4k and Supplementary Fig. 15b). The E325 in *Ps*PD forms a salt bridge with *Ps*PD R329 (corresponding to R328 in *Pa*PD), which causes a conformational shift of *Ps*PD R329, disrupting the salt bridge between *Ps*PD R329 and D459 in RBD (Fig. 4k and Supplementary Fig. 15b). The R33 in *Ps*PD (corresponding to H34 in *Pa*PD) forms new polar contacts with T514 and E551 of the HKU5 RBD (Fig. 4i and Supplementary Fig. 15c). Additionally, *Ps*PD displays a glycosylation pattern distinct from *Pa*PD. *Pa*PD carries multiple glycans (N53, N103, N215, N279, N386, N431, and N545), broadly distributed across its surface (Supplementary Fig. 14). In contrast, *Ps*PD has fewer sites (N52, N78, N322, and N546), which are more concentrated around the binding interface (Supplementary Fig. 14). Notably, N322 in *Ps*PD is glycosylated and forms additional contacts at the RBD–*Ps*PD interface (Fig. 4k and Supplementary Fig. 15d). These differences in glycosylation distribution likely contribute to the distinct binding modes and affinities observed for HKU5 RBD with *Pa*PD and *Ps*PD. Furthermore, compared to *Pa*PD, the overall structure of *Ps*PD is positioned slightly farther from the RBD, disrupting certain polar interactions, such as between *Pa*PD E37 and RBD K519 (Supplementary Fig. 15e).

To further investigate whether different HKU5 sub-clades affect RBD binding to *Ps*PD, we measured the KD values of the RBD of BtPa-BetaCoV/GD2013, HKU5-3, and other HKU5-related viruses for *Ps*PD, which were 137.8 ± 0.9 nM, 176.2 ± 1.4 nM, and 232.5 ± 2.0 nM, respectively (Supplementary Fig. 12). These differences suggest that mutations in different HKU5 strains may facilitate adaptation to ACE2 receptors from diverse species, potentially enabling cross-species transmission.

We analyzed the conservation of the RBD binding interface on ACE2 between bat and human. The results revealed substantial differences in the ^327^WRD^329^ and ^352^KND^354^ residues between bat and human ACE2, which may limit the potential for HKU5 to bind to human ACE2 (Fig. 4d). Although the spillover risk of HKU5 to human is currently low, continuous monitoring is warranted, especially in the context of viral evolution.

Together, these findings provide a comprehensive perspective on the structural, functional, and evolutionary characteristics of HKU5, enhancing our understanding of its receptor-binding specificity and potential for cross-species transmission.

## Discussion

This study provides critical insights into the molecular mechanisms underlying HKU5 S protein interaction with ACE2, shedding light on its structural adaptations, evolutionary dynamics, and the potential of cross-species transmission.

We identified two fatty acid molecules, oleic acid and palmitic acid, bound to the HKU5 S protein. These molecules stabilize the closed conformation of the S protein, likely modulating receptor accessibility. Similar to findings in other β-coronaviruses, including SARS-CoV-2^17^ and HCoV-OC43^40^, fatty acid binding occurs at conserved hydrophobic pockets within the RBD (Supplementary Table 1). Mutational analyses revealed that disruption of these interactions in HKU5 induces a structural shift toward a more relaxed conformation. This suggests that fatty acids play a crucial role in maintaining the closed state, potentially limiting premature receptor binding and aiding immune evasion. Notably, we identified a novel fatty acid binding pocket proximal to the RBM region in the HKU5 RBD. The cryo-EM map of the HKU5 RBD–*Pa*PD complex shows no density in this pocket, indicating that fatty acid binding may require the trimeric structure of the S protein. The regulatory mechanism aligns with prior studies proposing that fatty acid presence favors the “down” conformation of RBDs^41^. The widespread presence of fatty acids in coronaviral S proteins highlights their conserved role in regulating conformational dynamics. Efforts to design small-molecule inhibitors targeting these binding pockets have shown promise but require further structural optimization to enhance efficacy^42^. However, although cryo-EM and LC-MS identify oleic- and palmitic-acid binding at two sites on HKU5 S and pocket-disrupting mutations loosen the trimer, these substitutions did not cause a significant loss of pseudovirus entry. We infer that the fatty-acid pockets modulate the conformational equilibrium (stabilizing the closed state) without being obligatory for receptor engagement under our assay conditions.

Structural analysis revealed that the HKU5 RBD recognizes ACE2 through a unique mechanism distinct from other ACE2-binding coronaviruses. Unlike the RBM regions of SARS-CoV and SARS-CoV-2, which exhibit conserved ACE2-binding motifs^43^, HKU5 demonstrates an entirely novel interaction interface. These findings illustrate the evolutionary plasticity of coronaviruses and the convergent use of ACE2 by distinct lineages. Comparison of the available S protein structures from HKU5, HKU5-2, HKU5-19s, and MRCoV (Supplementary Fig. 16) shows a conserved closed conformation, while sequence diversification is clustered at the ACE2-binding interface. In particular, HKU5-2 carries substitutions within the RBM that may fine-tune receptor interactions, suggesting that adaptation primarily targets the binding interface rather than the overall spike architecture.

The broad tissue expression and high conservation of ACE2 across mammalian species provide a molecular basis for the cross-species transmission of coronaviruses^44,45^. Continuous adaptation of viral S proteins enables optimization of ACE2 binding, as exemplified by the progressive evolution of SARS-CoV-2 variants^46,47^. This interplay between receptor conservation and viral adaptability emphasizes the critical role of ACE2 in shaping coronavirus host range and transmission potential. Our analysis initially suggested that certain avian species, such as *Pitta sordida*, harbor ACE2 variants capable of binding the HKU5 RBD, raising the possibility of potential zoonotic reservoirs beyond mammals. In this study, we further substantiated this prediction by performing pseudovirus entry and cell–cell fusion assays, both of which demonstrated that cells stably expressing *Pitta sordida* ACE2 can indeed support HKU5 entry. These findings provide functional evidence that avian ACE2 orthologs are not only structurally compatible but also operationally competent to mediate HKU5 infection.

Traditionally, α- and β-coronaviruses are not thought to infect avian hosts, as birds are typically reservoirs for γ- and δ-coronaviruses. Our data therefore represent an intriguing exception and suggest that HKU5 has evolved a broader receptor usage than previously appreciated. This observation challenges the conventional view of strict host–virus associations and raises the possibility that certain avian species may, at least at the receptor level, serve as potential conduits for cross-class transmission. Nevertheless, we emphasize that our results are based on *in vitro* receptor reconstitution, and whether such interactions occur in natural settings will require live virus studies and ecological surveillance to assess their biological relevance.

In summary, our findings advance the understanding of how coronaviruses adapt to diverse hosts via ACE2 utilization, emphasizing the structural and functional diversity of this interaction. By elucidating the roles of fatty acid-mediated conformational regulation and ACE2 as a cross-species receptor, this study highlights the importance of monitoring HKU5 and related viruses for their zoonotic potential and public health implications.

## Methods

### Protein preparation

The extracellular domain (ECD) (1∼1293 a.a.) of the S protein of BatCoV-HKU5 (UniProt ID: A3EXD0) was cloned into the pCAG vector (Invitrogen) with a C-terminal T4 fibritin trimerization motif followed by a Flag tag and a 10×His tag. A “GSAS” mutation at residues 742 to 745 was introduced into the S protein to prevent cleavage by host. Double mutants of the S (S_DM1_) protein were generated with a standard two-step PCR-based strategy. The RBD domains of the S protein of different HKU5 sub-clades, including BatCoV-HKU5 (UniProt ID: A3EXD0), BatCoV-HKU5-3 (GenBank accession: ABN10893.1), BtPa-BetaCoV/GD2013 (GenBank accession: AIA62343.1) and BatCoV-HKU5-Realted (GenBank accession: QHA24687.1), were also cloned into the pCAG vector (Invitrogen) with an N-terminal signal peptide for secretion and a C-terminal 10×His tag. The PD domains of ACE2 of different species, including *Pipistrellus abramus* (UniProt ID: C7ECT9, *Pa*PD, 18∼615 a.a.), *Pitta sordida* (UniProt ID: A0A851FF99, *Ps*PD, 18∼617 a.a.) and *Homo sapiens* (UniProt ID: Q9BYF1, *Hs*PD, 25∼622 a.a.), were also cloned into the pCAG vector (Invitrogen) with an N-terminal signal peptide for secretion, followed by an N-terminal 2×StrepII tag and a C-terminal 10×His tag. All the plasmids used to transfect cells were prepared by GoldHi EndoFree Plasmid Maxi Kit (CWBIO). The purification processes of these proteins were the same as those described previously for the HKU1-B S protein^34^. Briefly, HEK293F mammalian cells (Thermo Fisher Scientific, Cat# A14527) were cultured in SMM293-TII medium (Sino Biological) at 37℃ and 5% CO_2_ in a Multitron-Pro shaker (Infors, 130 rpm). When the cell density reached 2.0×10^6^ cells/mL, the plasmid was transiently transfected into the cells. To transfect one liter of cell culture, approximately 1.5 mg of plasmid was premixed with 3 mg of polyethylenimine (PEI) (Yeasen Biotechnology) in 50 mL of fresh medium for 30 min before being added to the cell culture. After 70 h of overexpression, cells were removed by centrifugation at 4000×g for 15 min. The secreted proteins were purified by Ni-NTA Agarose (GE Healthcare). After loading twice, the Ni-NTA Agarose was washed with the wash buffer containing 25 mM Tris (pH 8.0), 150 mM NaCl, 50 mM imidazole. The protein was eluted with the elute buffer containing 25 mM Tris (pH 8.0), 150 mM NaCl, 500 mM imidazole. The eluate of RBDs or PDs was concentrated and subjected to size-exclusion chromatography (SEC, Superdex 200 Increase 10/300 GL, GE Healthcare) in buffer containing 25 mM Tris (pH 8.0), 150 mM NaCl. The eluent of S or S_DM1_ protein was subjected to SEC (Superose 6 Increase 10/300 GL, GE Healthcare) in buffer containing 25 mM Tris (pH 8.0), 150 mM NaCl. The HKU5 RBD was incubated with *Pa*PD or *Ps*PD at a molar ratio of about 5:1 for 1 h. Then the mixtures were subjected to SEC (Superose 6 Increase 10/300 GL, GE Healthcare) in buffer containing 25 mM Tris (pH 8.0), 150 mM NaCl. The peak fractions of SEC were collected and concentrated for cryo-EM analysis.

### Cryo-EM sample preparation and data acquisition

The protein samples were concentrated to ∼2.5 mg/mL and applied to the grids. Aliquots (3.5 μL) of the protein were placed on glow-discharged holey carbon grids (Quantifoil Au R1.2/1.3). The grids were blotted for 3.5 s and flash-frozen in liquid ethane cooled by liquid nitrogen using the Vitrobot Mark IV (Thermo Fisher Scientific). The prepared grids were transferred to a Krios (Thermo Fisher Scientific) operating at 300 kV equipped with a Gatan K3 detector and GIF Quantum energy filter. Movie stacks were automatically collected using EPU software (v3.11), with a slit width of 20 eV on the energy filter and a defocus range from -1.2 µm to -2.2 µm in super-resolution mode at a nominal magnification of 81,000×. Each stack was exposed for 2.56 s with an exposure time of 0.08 s per frame, resulting in a total of 32 frames per stack. The total dose rate was ∼50 e^-^/Å^2^ for each stack.

### Data processing

The movie stacks were motion corrected with MotionCor2^48^ and binned two fold, resulting in a pixel size of 1.087 Å/pixel. Meanwhile, dose weighting was performed^49^. After patch CTF estimation, particles were automatically picked according to the templates generated from an initial 2D classification of manually picked particles by cryoSPARC (v4.6.2)^50^. After 2D classification, the particles with clear features were selected and subjected to *ab-initio* reconstruction to obtain the initial models, then multi-hetero refinements without symmetry were performed to selected good particles using cryoSPARC (v4.6.2)^50^. The selected particles were subjected to non-uniform refinement, local CTF refinement or global CTF refinement, and local refinement, resulting in 3D reconstruction for the whole structure. The resolution was estimated with the gold-standard Fourier shell correlation 0.143 criterion^51^ with high-resolution noise substitution^52^. Minor variations in processing were applied for different samples, as detailed in Supplementary Figures 1, 4, 7, and 13. All of the map figures were generated in ChimeraX (v1.3)^53^ or Chimera (v1.16)^54^.

### Model building and structure refinement

The predicted atomic models generated by Alphafold 3^55^ were used as templates and flexibly fitted into the cryo-EM maps using molecular dynamics in VMD (v1.9.3)^56^, followed by manual adjustment with Coot (v0.8.9.2)^57^ to obtain the atomic models. Each residue was manually checked with the chemical properties taken into consideration during model building. Several segments, whose corresponding densities were invisible, were not modeled. Specifically, in both the HKU5 S protein and the S_DM1_ protein, the 739∼750 a.a. at the S1/S2 cleavage site could not be modeled. In addition, 686∼700 a.a. within SD2 of the HKU5 S protein were not built. Structural refinement was performed in Phenix (v1.14)^58^ with secondary structure and geometry restraints to prevent overfitting (Supplementary Table 2).

### Generation of stable cell lines

Full-length ACE2 from three species, *Pipistrellus abramus*, *Pitta sordida*, and *Homo sapiens*, were cloned into the lentiCRISPR v2 vector^59^ with a spleen focus-forming virus (SFFV) promoter, followed by a FLAG tag and linked via a P2A ribosome-skipping site (ATNFSLLKQAGDVEENPGP) to an mCherry reporter at the C terminus. Mutants of P.abr ACE2 were cloned using the same strategy. The lentiviral transfer plasmids encoding full-length ACE2, together with the packaging vectors psPAX2 and pMD2.G, were co-transfected into Lenti-X 293T cells (Takara Bio, Cat# 632180). Cells were maintained in D10 medium (DMEM supplemented with 10% (v/v) FBS, 100 U/mL penicillin G, and 100 μg/mL streptomycin) at 37 °C in a water-saturated atmosphere containing 5% CO_2_. Three days after transfection, the supernatants were harvested, clarified by centrifugation at 1,000×g, and filtered through 0.45 μm membranes (Beyotime). To concentrate lentivirus, protamine sulfate (80 μg/mL, Macklin) and chondroitin sulfate C (80 μg/mL, Macklin) were added as described previously^60^. Next, Lenti-X 293T or Caco-2 (ATCC, Cat# HTB-37) cells (1×10⁶ per well in 1 mL D10) supplemented with 5 μg/mL polybrene (Beyotime) were transduced with 1 mL concentrated lentivirus. Cells exhibiting comparable mCherry fluorescence intensity were sorted on a MA900 Multi-Application Cell Sorter (Sony). Debris was excluded using FSC-A versus SSC-A, singlets were selected using FSC-A versus FSC-H, and mCherry-positive cells were gated in the 561-nm channel (610/20-nm filter). The sort gate was centered on the main mCherry-positive population to ensure comparable expression across lines. Sorted cells were used for subsequent experiments.

### Pseudovirus production and entry assays

Full-length HKU5 (UniProt ID: A3EXD0) S proteins, with or without mutations, were cloned into the pCAG vector (Invitrogen) with a C-terminal truncation (Δ13) followed by a 10×His tag. Pseudoviruses (PSVs) were generated using the HIV-1 backbone plasmid pNL4-3.Luc.R⁻E⁻^61^, which encodes a luciferase reporter but lacks functional Env. Briefly, Lenti-X 293T cells were seeded into 10-cm dishes and co-transfected with pNL4-3.Luc.R⁻E⁻ (20 µg) and HKU5 S plasmid (10 µg) using PEI (90 µg). Supernatants were harvested 48 h post-transfection, clarified by centrifugation at 12,000×g for 10 min at 4 °C, aliquoted, and stored at −80 °C.

For entry assays, Lenti-X 293T cells stably or transiently expressing indicated ACE2 orthologs or mutants were seeded in 96-well plates (5×10⁴ cells/well) and infected with PSVs in the presence of polybrene (8 μg/mL, Beyotime) for 18 h. Before inoculation, PSVs produced in Opti-MEM (Gibco) were treated with TPCK-trypsin (25 μg/mL, Sigma-Aldrich, T8802) at room temperature for 10 min, and proteolytic activity was neutralized by FBS. Next, cells were replenished with D10 medium, cultured for an additional 48 h, and then lysed with Cell Culture Lysis 5× Reagent (Promega, diluted 1:5 in deionized water) at room temperature for 30 min. Luciferase activity was then measured with the Luciferase Assay System (Promega, E1501) using a Varioskan LUX multimode microplate reader (Thermo Fisher Scientific). All assays were performed in triplicate, and data were analyzed and plotted using GraphPad Prism (v10.1.2).

### Western blot

Western blot was performed to examine the proteolytic processing and incorporation efficiency of WT and mutant HKU5 S proteins. The PSV-containing supernatants were concentrated using protamine sulfate (80 μg/mL, Macklin) and chondroitin sulfate C (80 μg/mL, Macklin). Viral pellets were resuspended in Cell Culture Lysis 5× Reagent (Promega, diluted 1:5 in deionized water) and incubated at room temperature for 10 min, followed by mixing with one-fifth of the volume of 5×SDS loading buffer. Proteins were resolved by SDS–PAGE and probed using anti His-Tag mouse monoclonal antibody (CWBIO, CW0286M; 1:2000) followed by goat anti-mouse IgG secondary antibody (CWBIO, CW0102; 1:5000). Blots were washed three times with TBST and visualized using the BeyoECL Plus (Beyotime, P0018S) with Amersham Imager 680 (Cytiva).

### Bio-layer interferometry (BLI)

BLI experiments were performed using the Octet system (ForteBio) to characterize the protein interactions. StrepII-10×His-tagged *Pa*PD and *Ps*PD proteins, along with 10×His-RBD proteins from four HKU5 strains, were purified separately. At room temperature, *Pa*PD proteins were immobilized onto equilibrated streptavidin (SA) biosensors. Subsequently, the biosensors were exposed to solutions containing various concentration of RBD proteins (0 nM, 12.5 nM, 25 nM, 50 nM, 100 nM, 200 nM, 400 nM, and 800 nM), with binding kinetics monitored in real-time. Following the association phase, the biosensors were immersed in the assay buffer to monitor dissociation kinetics in real-time. Sensorgrams were analyzed using Octet Analysis Studio (v13.0) to determine the binding affinity constants between the PD and RBD proteins. This process was repeated to evaluate the binding affinities between the two PD proteins and the four RBD proteins. For the binding affinity measurements between *Ps*PD and different RBDs, the RBD concentration gradient was set at 0 µM, 0.04 µM, 0.08 µM, 0.16 µM, 0.32 µM, 0.64 µM, 1.28 µM, and 2.56 µM.

### Tripartite split-fluorescence system

We designed a tripartite split-fluorescence system containing plasmids A, B, and C. Plasmid A encoded the first nine β-strands (1∼196 a.a.) of sfGFP, followed by a stop codon and an internal ribosome entry site 2 (iRES2) sequence to mediate the subsequent translation of iRFP702. For plasmid B, the tenth β-strand of sfGFP (197∼216 a.a.) was inserted at the N-terminus and fused to the RBD via a linker (DVGGGGSEGGGSGGPGSGGEGSAGGGSAGGGS). An iRES2 sequence and mCherry sequence were then inserted downstream. For plasmid C, the PD was inserted at the N-terminus and fused to the eleventh β-strand of sfGFP (217∼238 a.a.) via a linker (GSGAGGSPGGGSGGSGSSASGGSTSGGGSGGGS), followed by iRES2 and EBFP2 sequences. The designed sequences were synthesized by SynbioB and cloned into the mammalian expression vector pCAG.

### Flow cytometry sorting and statistical analysis

HEK293F cells were cultured in SMM293-TII medium (Sino Biological) at 37°C with 5% CO_2_ under shaking conditions (120 rpm). Plasmid A, plasmid B expressing WT or mutant RBD, and plasmid C expressing WT or mutant PD were mixed at a 1:1:1 ratio (5 µg each) and transfected into cells (2×10⁶ cells per 5 mL flask) using PEI at a 2:1 DNA-to-PEI mass ratio. After 48 h, cells were harvested by centrifugation at 200×g for 2 min and washed twice with 2 mL PBS (Gibco) at room temperature. The final cell pellet was resuspended in 1ml PBS for flow cytometry. Flow cytometry was performed on a BD FACSymphony^TM^ A3 (BD Biosciences). Single-cell populations were gated based on forward scatter area (FSC-A), side scatter area (SSC-A), and forward scatter height (FSC-H) parameters. Cells co-expressing plasmids A, B, and C, identified by simultaneous positivity in the BV421 (405 nm laser, 450/50 nm filter), BV711 (405 nm laser, 710/50 nm filter), and PE-CF594 (561 nm laser, 610/20 nm filter) fluorescence channels, were analyzed for sfGFP fluorescence intensity in the FITC channel (488 nm laser, 530/30 nm filter). The median fluorescence intensity of the gated population was calculated using FlowJo (v10.6.2). All experiments were performed in triplicate.

### Immunofluorescence staining

Immunofluorescence assays were performed to assess the expression of ACE2 orthologs and mutants fused with FLAG tags. Transfected cells were fixed and permeabilized with 100% methanol for 10 min at room temperature, incubated with mouse anti-DDDDK-tag mAb (ABclonal, AE005; 1:200) diluted in PBS containing 1% BSA for 1 h at 37 °C, washed with HBSS, and then incubated with ABflo® 647-conjugated goat anti-mouse IgG (ABclonal, AS059; 1:150) for 1 h at 37 °C in the dark. Nuclei were stained with Hoechst 33342 (1:100 dilution in HBSS) for 30 min at 37 °C. Images were acquired using a CSU-W1 SoRa spinning disk confocal system (Nikon) and processed with Fiji (ImageJ v1.54f).

### Cell–cell fusion assay

Cell–cell fusion was performed in Caco-2 cells stably expressing ACE2 orthologs using a dual-split protein (DSP) reporter system. To evaluate membrane fusion mediated by S glycoprotein-ACE2, group A cells were transfected with HKU5 S and rLucN(1–155)-sfGFP1–7(1–157) plasmids, while group B cells were transfected with HKU5 S and sfGFP8–11(158–231)-rLuc(156–311). After 24 h, cells were trypsinized, mixed, and seeded into 96-well plates (8×10⁴ cells/well). Following another 24 h of growth, cells were washed with DMEM and incubated with or without TPCK-trypsin (Sigma-Aldrich, T8802) for 10 min at room temperature, followed by immediate replenishment with D10 medium to neutralize trypsin. After 12 h, nuclei were stained with Hoechst 33342 (1:100 dilution in HBSS) for 30 min at 37 °C, and syncytia formation with green fluorescence was visualized using a CSU-W1 SoRa spinning disk confocal system (Nikon) and processed with Fiji (ImageJ v1.54f).

## Data availability

The atomic coordinates and cryo-EM maps generated in this study have been deposited in the Protein Data Bank (http://www.rcsb.org) and the Electron Microscopy Data Bank (https://www.ebi.ac.uk/pdbe/emdb/) under the following accession codes (PDB: 9KR8, 9KR9, 9KRA, 9KRB; EMDB: EMD-62522, EMD-62523, EMD-62524, EMD-62525, EMD-64945).

All other data supporting the findings of this study are available within the article and the Supplementary Information. All materials used in this study are available from the corresponding authors upon reasonable request. Source data are provided with this paper.

## Acknowledgements

This work is supported by National Natural Science Foundation of China (82241081), the Key Regional Research and Development Program (2023CSJZN0600) from Ministry of Science and Technology of China, and State Key Laboratory of Gene Expression. We thank the staff of the cryo-EM facility, the high-performance computing center, the mass spectrometry and metabolomics core facility, the flow cytometry core facility, the protein characterization and crystallography facility, and the microscopy imaging platform of Westlake University for technical assistance, advice, and support. We also thank Dr. Xu Li and Dr. Peihan Wu for providing materials and technical assistance for pseudovirus preparation.

## Author contributions

Q.Z. and L.X. conceived the project. Y.Z., L.X., Y.L., W.G., and D.L. performed the experiments. Y.Z. and L.X. analyzed the PDB structures and performed data visualization. All authors contributed to data analysis. Q.Z., L.X., and Y.Z. wrote the manuscript. Q.Z. supervised the project.

## References

1 Fouchier, R. A. et al. Aetiology: Koch’s postulates fulfilled for SARS virus. Nature 423, 240, doi:10.1038/423240a (2003).

2 Drosten, C. et al. Identification of a novel coronavirus in patients with severe acute respiratory syndrome. N Engl J Med 348, 1967–1976, doi:10.1056/NEJMoa030747 (2003).

3 Wu, F. et al. A new coronavirus associated with human respiratory disease in China. Nature 579, 265–269, doi:10.1038/s41586-020-2008-3 (2020).

4 Zhou, P. et al. A pneumonia outbreak associated with a new coronavirus of probable bat origin. Nature 579, 270–273, doi:10.1038/s41586-020-2012-7 (2020).

5 Ruiz-Aravena, M. et al. Ecology, evolution and spillover of coronaviruses from bats. Nature Reviews Microbiology 20, 299–314, doi:10.1038/s41579-021-00652-2 (2022).

6 Mackenzie, J. S., Childs, J. E., Field, H. E., Wang, L. F. & Breed, A. C. The Role of Bats as Reservoir Hosts of Emerging Neuroviruses. (Neurotropic Viral Infections. 2016 Apr 8:403–54. doi: 10.1007/978-3-319-33189-8_12.).

7 Letko, M., Seifert, S. N., Olival, K. J., Plowright, R. K. & Munster, V. J. Bat-borne virus diversity, spillover and emergence. Nature Reviews Microbiology 18, 461–471, doi:10.1038/s41579-020-0394-z (2020).

8 Jackson, C. B., Farzan, M., Chen, B. & Choe, H. Mechanisms of SARS-CoV-2 entry into cells. Nat Rev Mol Cell Biol 23, 3–20, doi:10.1038/s41580-021-00418-x (2022).

9 Benton, D. J. et al. Receptor binding and priming of the spike protein of SARS-CoV-2 for membrane fusion. Nature 588, 327–330, doi:10.1038/s41586-020-2772-0 (2020).

10 Gui, M. et al. Cryo-electron microscopy structures of the SARS-CoV spike glycoprotein reveal a prerequisite conformational state for receptor binding. Cell Res 27, 119–129, doi:10.1038/cr.2016.152 (2017).

11 Wrapp, D. et al. Cryo-EM structure of the 2019-nCoV spike in the prefusion conformation. Science 367, 1260–1263, doi:10.1126/science.abb2507 (2020).

12 Ke, Z. et al. Structures and distributions of SARS-CoV-2 spike proteins on intact virions. Nature 588, 498–502, doi:10.1038/s41586-020-2665-2 (2020).

13 Nguyen, L. et al. Sialic acid-containing glycolipids mediate binding and viral entry of SARS-CoV-2. Nature Chemical Biology 18, 81–90, doi:10.1038/s41589-021-00924-1 (2022).

14 Pronker, M. F. et al. Sialoglycan binding triggers spike opening in a human coronavirus. Nature 624, 201–206, doi:10.1038/s41586-023-06599-z (2023).

15 Hulswit, R. J. G. et al. Human coronaviruses OC43 and HKU1 bind to 9-O-acetylated sialic acids via a conserved receptor-binding site in spike protein domain A. Proc Natl Acad Sci U S A 116, 2681–2690, doi:10.1073/pnas.1809667116 (2019).

16 Qing, E., Hantak, M., Perlman, S. & Gallagher, T. Distinct Roles for Sialoside and Protein Receptors in Coronavirus Infection. mBio 11, doi:10.1128/mBio.02764-19 (2020).

17 Toelzer, C. et al. Free fatty acid binding pocket in the locked structure of SARS-CoV-2 spike protein. Science 370, 725–730, doi:10.1126/science.abd3255 (2020).

18 de Groot, R. J. et al. Middle East respiratory syndrome coronavirus (MERS-CoV): announcement of the Coronavirus Study Group. J Virol 87, 7790–7792, doi:10.1128/jvi.01244-13 (2013).

19 Lau, S. K. et al. Genetic characterization of Betacoronavirus lineage C viruses in bats reveals marked sequence divergence in the spike protein of pipistrellus bat coronavirus HKU5 in Japanese pipistrelle: implications for the origin of the novel Middle East respiratory syndrome coronavirus. J Virol 87, 8638–8650, doi:10.1128/jvi.01055-13 (2013).

20 Woo, P. C., Lau, S. K., Li, K. S., Tsang, A. K. & Yuen, K. Y. Genetic relatedness of the novel human group C betacoronavirus to Tylonycteris bat coronavirus HKU4 and Pipistrellus bat coronavirus HKU5. Emerg Microbes Infect 1, e35, doi:10.1038/emi.2012.45 (2012).

21 Woo, P. C. Y. et al. Molecular diversity of coronaviruses in bats. Virology 351, 180–187, 10.1016/j.virol.2006.02.041 (2006).

22 Chen, J. et al. Bat-infecting merbecovirus HKU5-CoV lineage 2 can use human ACE2 as a cell entry receptor. Cell 188, 1729–1742.e1716, doi:10.1016/j.cell.2025.01.042 (2025).

23 Zhao, J. et al. Farmed fur animals harbour viruses with zoonotic spillover potential. Nature 634, 228–233, doi:10.1038/s41586-024-07901-3 (2024).

24 Park, Y. J. et al. Molecular basis of convergent evolution of ACE2 receptor utilization among HKU5 coronaviruses. Cell 188, 1711–1728.e1721, doi:10.1016/j.cell.2024.12.032 (2025).

25 Catanzaro, N. J. et al. ACE2 from Pipistrellus abramus bats is a receptor for HKU5 coronaviruses. Nat Commun 16, 4932, doi:10.1038/s41467-025-60286-3 (2025).

26 Li, W. et al. Angiotensin-converting enzyme 2 is a functional receptor for the SARS coronavirus. Nature 426, 450–454, doi:10.1038/nature02145 (2003).

27 Hoffmann, M. et al. SARS-CoV-2 Cell Entry Depends on ACE2 and TMPRSS2 and Is Blocked by a Clinically Proven Protease Inhibitor. Cell 181, 271–280.e278, doi:10.1016/j.cell.2020.02.052 (2020).

28 Hofmann, H. et al. Human coronavirus NL63 employs the severe acute respiratory syndrome coronavirus receptor for cellular entry. Proc Natl Acad Sci U S A 102, 7988–7993, doi:10.1073/pnas.0409465102 (2005).

29 Ma, C. B. et al. Multiple independent acquisitions of ACE2 usage in MERS-related coronaviruses. Cell 188, 1693–1710.e1618, doi:10.1016/j.cell.2024.12.031 (2025).

30 Jiang, S. & Wu, F. Global surveillance and countermeasures for ACE2-using MERS-related coronaviruses with spillover risk. Cell 188, 1465–1468, doi:10.1016/j.cell.2025.02.004 (2025).

31 Liu, C. et al. HKU25 clade MERS-related coronaviruses use ACE2 as a functional receptor. Nature Microbiology 10, 2860–2874, doi:10.1038/s41564-025-02152-y (2025).

32 Medina-Enríquez, M. M. et al. ACE2: the molecular doorway to SARS-CoV-2. Cell Biosci 10, 148, doi:10.1186/s13578-020-00519-8 (2020).

33 Yuan, Y. et al. Cryo-EM structures of MERS-CoV and SARS-CoV spike glycoproteins reveal the dynamic receptor binding domains. Nat Commun 8, 15092, doi:10.1038/ncomms15092 (2017).

34 Xia, L., Zhang, Y. & Zhou, Q. Structural basis for the recognition of HCoV-HKU1 by human TMPRSS2. Cell Res 34, 526–529, doi:10.1038/s41422-024-00958-9 (2024).

35 Staufer, O. et al. Synthetic virions reveal fatty acid-coupled adaptive immunogenicity of SARS-CoV-2 spike glycoprotein. Nature Communications 13, 868, doi:10.1038/s41467-022-28446-x (2022).

36 Yan, R. et al. Structural basis for the different states of the spike protein of SARS-CoV-2 in complex with ACE2. Cell Res 31, 717–719, doi:10.1038/s41422-021-00490-0 (2021).

37 Xiong, Q. et al. Close relatives of MERS-CoV in bats use ACE2 as their functional receptors. Nature 612, 748–757, doi:10.1038/s41586-022-05513-3 (2022).

38 Starr, T. N. et al. ACE2 binding is an ancestral and evolvable trait of sarbecoviruses. Nature 603, 913–918, doi:10.1038/s41586-022-04464-z (2022).

39 Chakrabarti, S. et al. Touch sensation requires the mechanically gated ion channel ELKIN1. Science 383, 992–998, doi:10.1126/science.adl0495 (2024).

40 Bangaru, S. et al. Structural mapping of antibody landscapes to human betacoronavirus spike proteins. Sci Adv 8, eabn2911, doi:10.1126/sciadv.abn2911 (2022).

41 Hills, F. R. et al. Variation in structural motifs within SARS-related coronavirus spike proteins. PLoS Pathog 20, e1012158, doi:10.1371/journal.ppat.1012158 (2024).

42 Wang, Q. et al. In Silico Discovery of Small Molecule Modulators Targeting the Achilles’ Heel of SARS-CoV-2 Spike Protein. ACS Cent Sci 9, 252–265, doi:10.1021/acscentsci.2c01190 (2023).

43 Yan, R. et al. Structural basis for the recognition of SARS-CoV-2 by full-length human ACE2. Science 367, 1444–1448, doi:10.1126/science.abb2762 (2020).

44 Damas, J. et al. Broad host range of SARS-CoV-2 predicted by comparative and structural analysis of ACE2 in vertebrates. Proc Natl Acad Sci U S A 117, 22311–22322, doi:10.1073/pnas.2010146117 (2020).

45 Hamming, I. et al. Tissue distribution of ACE2 protein, the functional receptor for SARS coronavirus. A first step in understanding SARS pathogenesis. J Pathol 203, 631–637, doi:10.1002/path.1570 (2004).

46 Li, L. et al. Structural basis of human ACE2 higher binding affinity to currently circulating Omicron SARS-CoV-2 sub-variants BA.2 and BA.1.1. Cell 185, 2952–2960.e2910, doi:10.1016/j.cell.2022.06.023 (2022).

47 Feng, L. et al. Structural and molecular basis of the epistasis effect in enhanced affinity between SARS-CoV-2 KP.3 and ACE2. Cell Discov 10, 123, doi:10.1038/s41421-024-00752-2 (2024).

48 Zheng, S. Q. et al. MotionCor2: anisotropic correction of beam-induced motion for improved cryo-electron microscopy. Nat Methods 14, 331–332, doi:10.1038/nmeth.4193 (2017).

49 Grant, T. & Grigorieff, N. Measuring the optimal exposure for single particle cryo-EM using a 2.6 Å reconstruction of rotavirus VP6. Elife 4, e06980, doi:10.7554/eLife.06980 (2015).

50 Punjani, A., Rubinstein, J. L., Fleet, D. J. & Brubaker, M. A. cryoSPARC: algorithms for rapid unsupervised cryo-EM structure determination. Nat Methods 14, 290–296, doi:10.1038/nmeth.4169 (2017).

51 Rosenthal, P. B. & Henderson, R. Optimal determination of particle orientation, absolute hand, and contrast loss in single-particle electron cryomicroscopy. J Mol Biol 333, 721–745, doi:10.1016/j.jmb.2003.07.013 (2003).

52 Chen, S. et al. High-resolution noise substitution to measure overfitting and validate resolution in 3D structure determination by single particle electron cryomicroscopy. Ultramicroscopy 135, 24–35, doi:10.1016/j.ultramic.2013.06.004 (2013).

53 Pettersen, E. F. et al. UCSF ChimeraX: Structure visualization for researchers, educators, and developers. Protein Sci 30, 70–82, doi:10.1002/pro.3943 (2021).

54 Pettersen, E. F. et al. UCSF Chimera--a visualization system for exploratory research and analysis. J Comput Chem 25, 1605–1612, doi:10.1002/jcc.20084 (2004).

55 Abramson, J. et al. Accurate structure prediction of biomolecular interactions with AlphaFold 3. Nature 630, 493–500, doi:10.1038/s41586-024-07487-w (2024).

56 Trabuco, L. G., Villa, E., Mitra, K., Frank, J. & Schulten, K. Flexible fitting of atomic structures into electron microscopy maps using molecular dynamics. Structure 16, 673–683, doi:10.1016/j.str.2008.03.005 (2008).

57 Emsley, P., Lohkamp, B., Scott, W. G. & Cowtan, K. Features and development of Coot. *Acta Crystallogr D Biol Crystallogr* **66**, 486–501, doi:10.1107/s0907444910007493 (2010).

58 Adams, P. D. et al. PHENIX: a comprehensive Python-based system for macromolecular structure solution. Acta Crystallogr D Biol Crystallogr 66, 213–221, doi:10.1107/s0907444909052925 (2010).

59 Sanjana, N. E., Shalem, O. & Zhang, F. Improved vectors and genome-wide libraries for CRISPR screening. Nat Methods 11, 783–784, doi:10.1038/nmeth.3047 (2014).

60 Lee, J. Y. & Lee, H. H. A new chemical complex can rapidly concentrate lentivirus and significantly enhance gene transduction. Cytotechnology 70, 193–201, doi:10.1007/s10616-017-0133-0 (2018).

61 Connor, R. I., Chen, B. K., Choe, S. & Landau, N. R. Vpr is required for efficient replication of human immunodeficiency virus type-1 in mononuclear phagocytes. Virology 206, 935–944, doi:10.1006/viro.1995.1016 (1995).

