## Supplementary information for "Molecular Insights into Species-Specific ACE2 Recognition of Coronavirus HKU5"

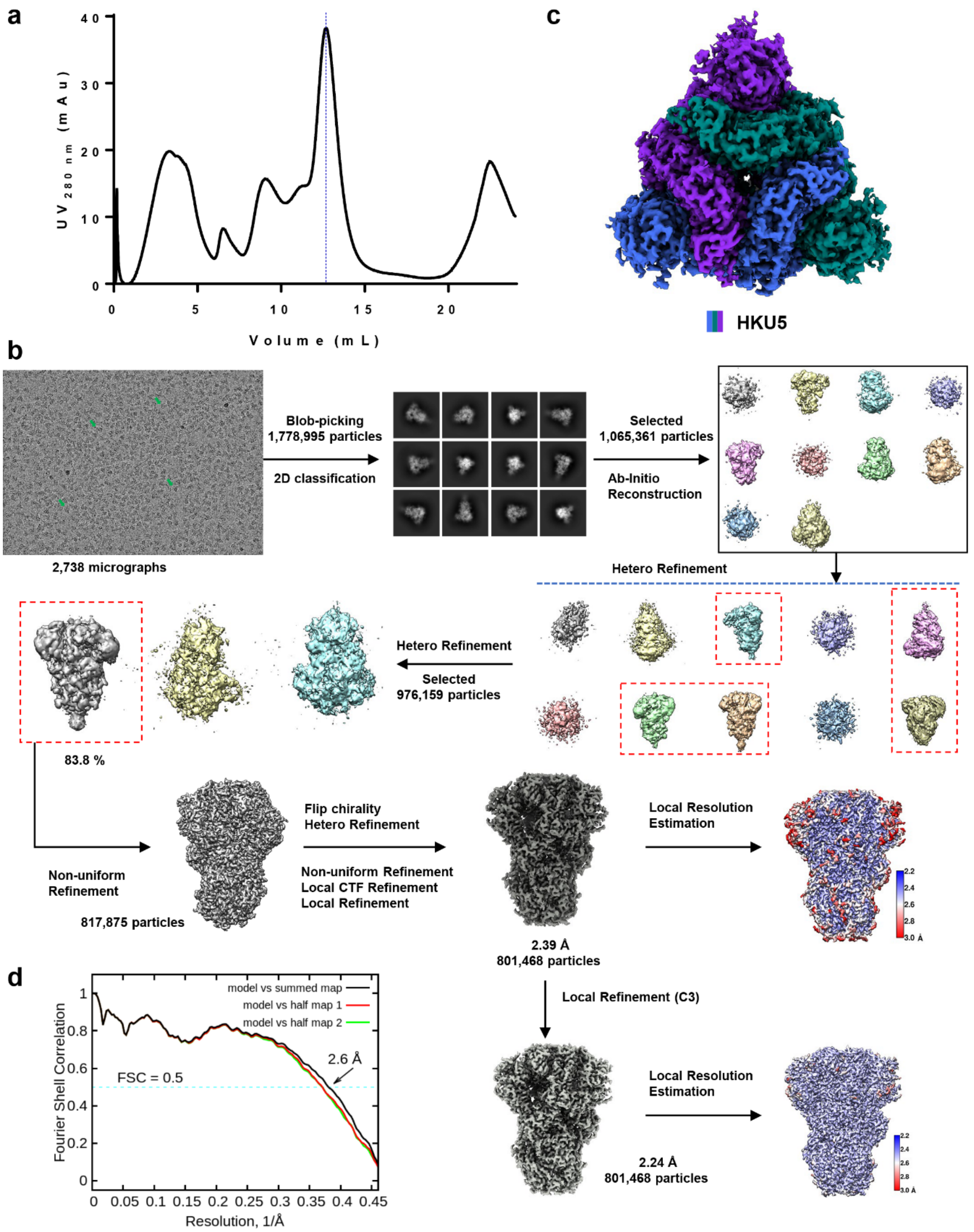


**Supplementary Fig. 1. Cryo-EM data processing and analysis for the HKU5 S protein.**

**a,** Gel filtration elution profile of HKU5 S protein. **b,** Flowchart of cryo-EM data processing, see the "Data Processing" section in Methods for details. **c,** Cryo-EM map shown in top view at a contour level of 4 σ of the HKU5 S protein, with protomers displayed in marine blue, purple, and forest green, respectively. **d,** FSC curve of the refined model of HKU5 S protein versus the overall map that it is refined against (black); of the model refined against the first half map versus the same map (red); and of the model refined against the first half map versus the second half map (green). The small difference between the red and green curves indicates that the refinement of the atomic coordinates did not suffer from overfitting.

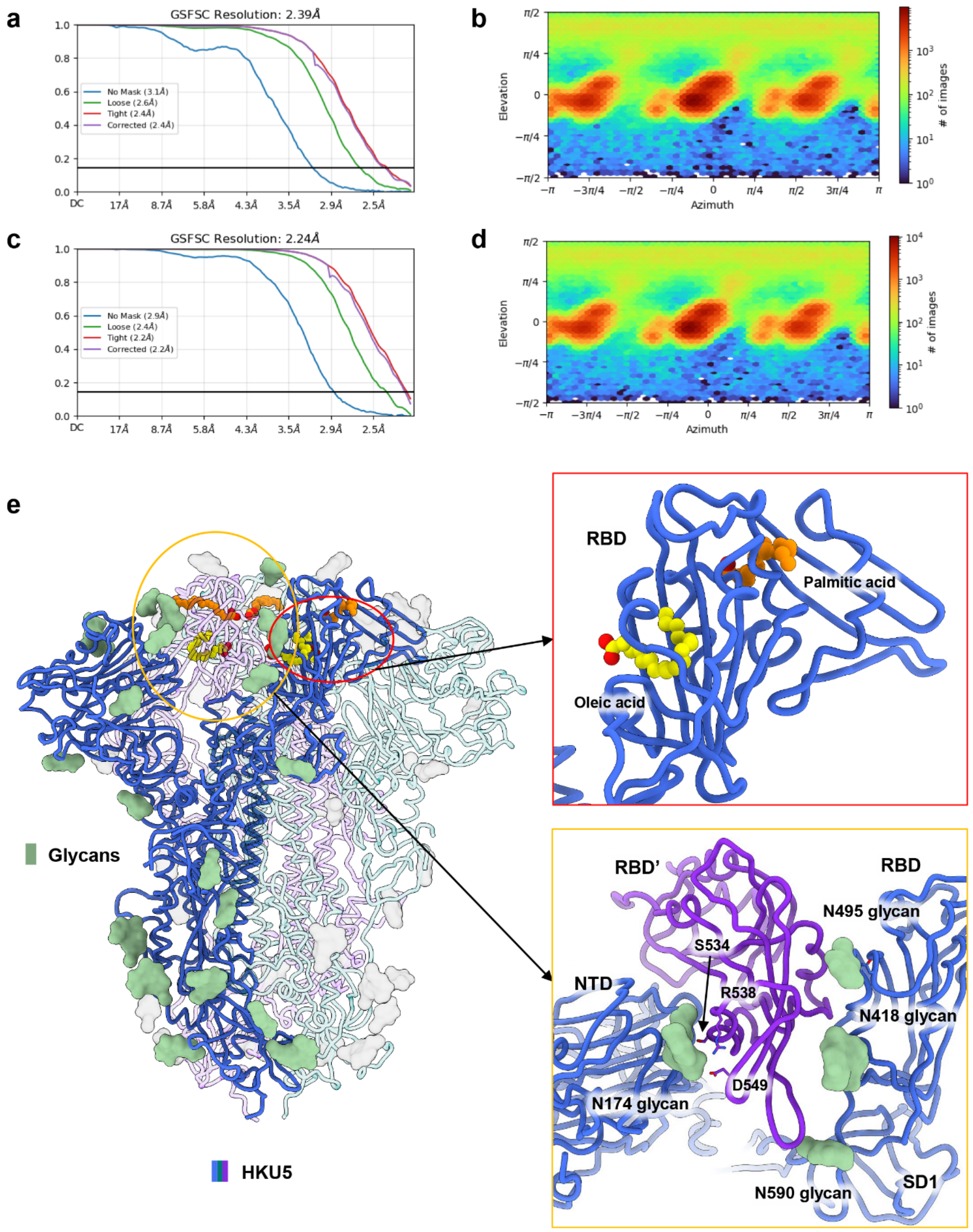


**Supplementary Fig. 2. Analysis for the HKU5 S protein.**

**a-b,** Gold standard FSC curve and euler angle distribution in C1 symmetry of the HKU5 S protein is estimated by cryoSPARC. **c-d,** Gold standard FSC curve and euler angle distribution in C3 symmetry of the HKU5 S protein is estimated by cryoSPARC. **e,** Overall structure of the HKU5 S protein, highlighting one protomer shown in marine blue. N-linked glycans are displayed as green surfaces, oleic acid as yellow spheres, and palmitic acid as orange spheres. The red inset panel presents an enlarged view of the fatty acid–binding sites within the HKU5 S protein. The yellow inset panel shows structural details of the RBD surrounded by multiple N-linked glycans (N174, N418, N495, and N590), revealing that the RBD is positioned in the "down" conformation and extensively shielded by glycans.

a

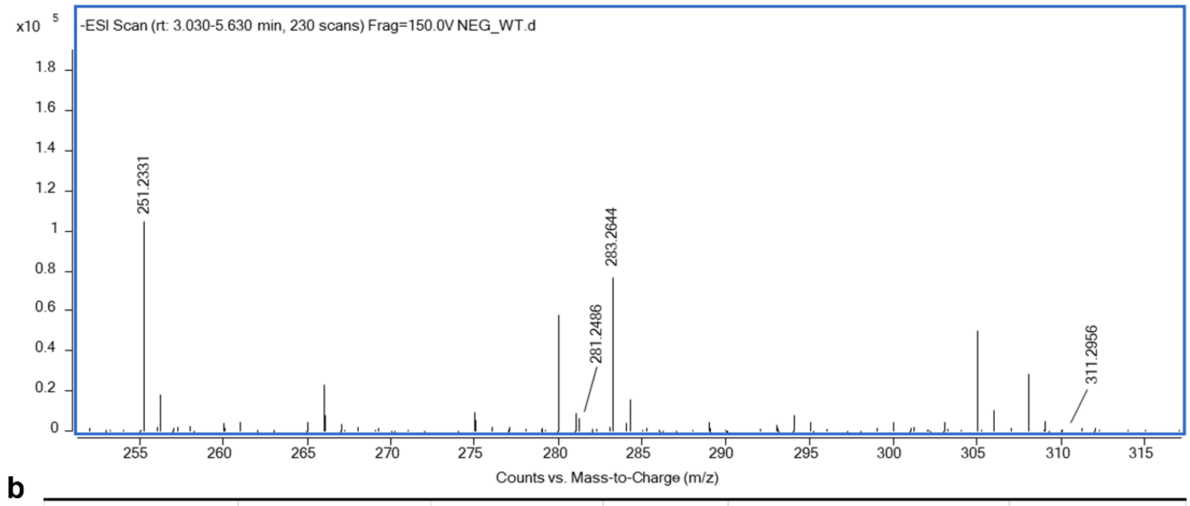


b

| Compound Name | Formula | Mass | RT | NEG_WT（Area） | SNR |
|---|---|---|---|---|---|
| FA 18:0 | C18 H36 O2 | 284.2716 | 4.37 | 11196418.18 | 252.2 |
| FA 16:0 | C16 H32 O2 | 256.2404 | 3.44 | 15968231.22 | 15874.9 |
| FA 18:1 | C18 H34 O2 | 282.2556 | 3.64 | 674911.7934 | 6176.9 |
| FA 20:0 | C20 H40 O2 | 312.3026 | 5.19 | 215788.3878 | 18.3 |

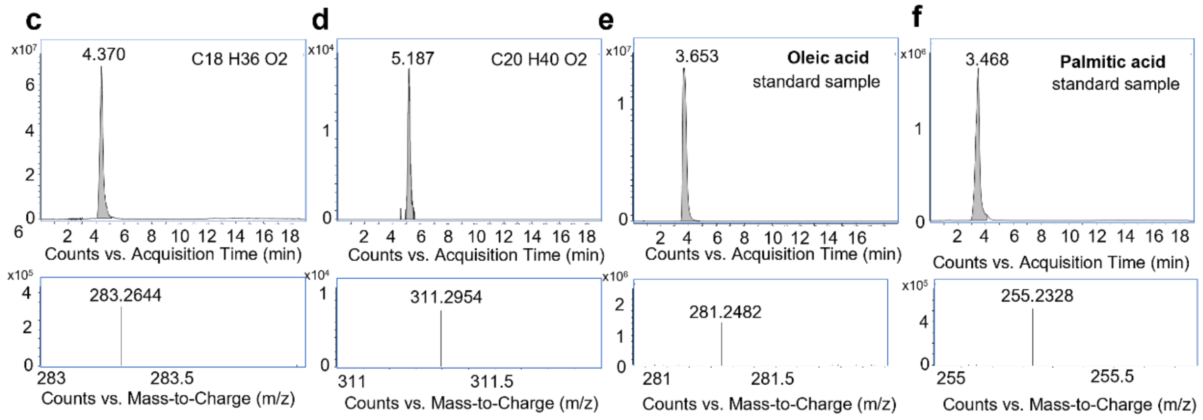


**Supplementary Fig. 3. Mass spectrometry characterization of the HKU5 S protein.**

**a,** The mass spectrometry profile of the HKU5 S protein. The x-axis represents the m/z ratio, which is the mass-to-charge ratio of the ions. The y-axis represents the ion intensity, reflecting the abundance of the ions in the sample. Each peak in the mass spectrum corresponds to a specific ion. **b,** Information on the identified compounds. RT: retention time, NEG_WT: negative ion peak area, SNR: signal-to-noise ratio. The NEG_WT is proportional to the compound concentration. **c-d,** Mass spectra of octadecanoic acid (FA 18:0) and eicosanoic acid (FA 20:0), respectively. **e-f,** Standard mass spectra of oleic acid and palmitic acid, respectively. These spectra serve as references for the qualitative identification of compounds in the sample.

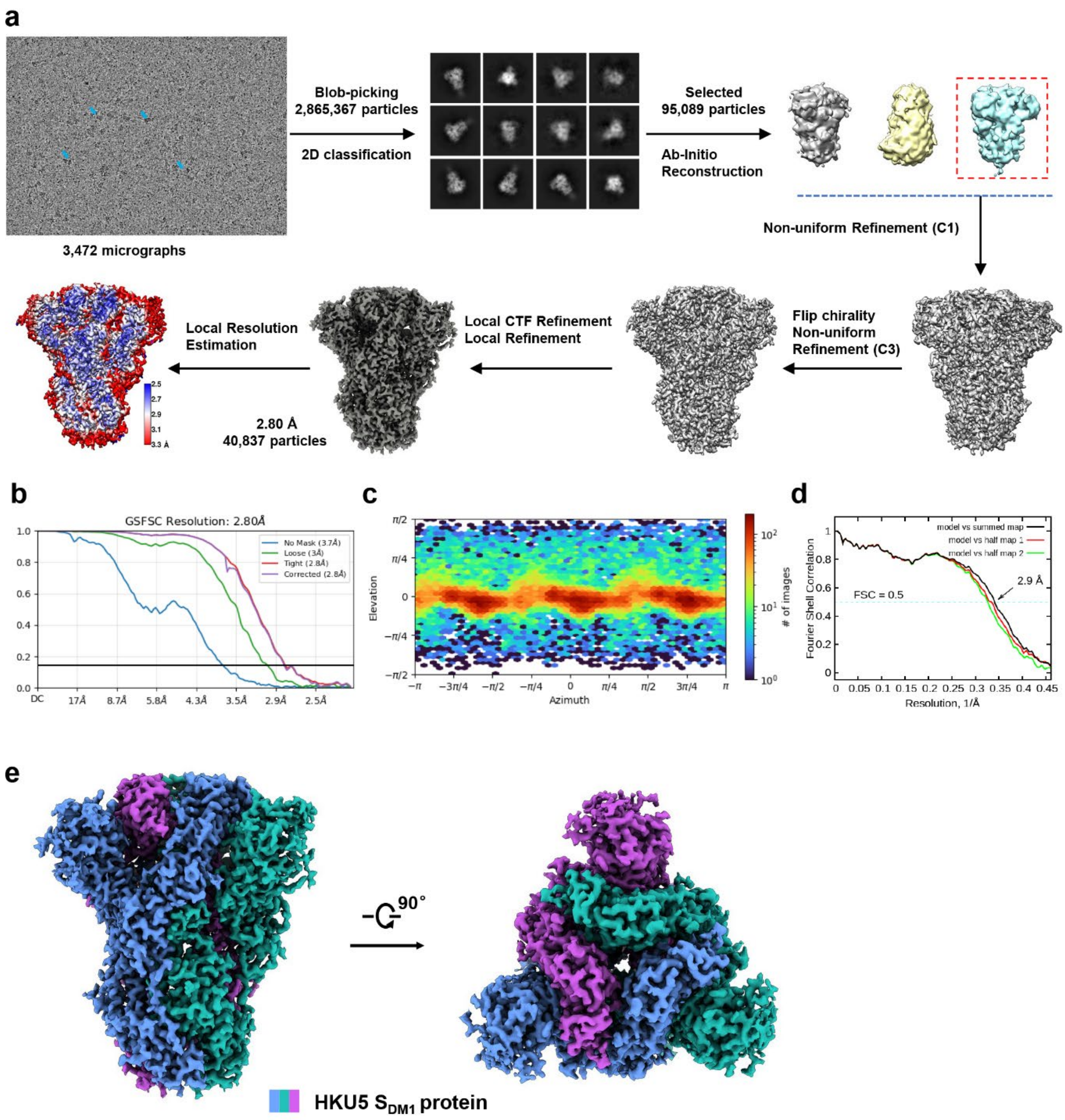


**Supplementary Fig. 4. Cryo-EM data processing and analysis for the HKU5 $S_{DM1}$ protein.**

**a,** Flowchart of cryo-EM data processing, see the “Data Processing” section in Methodsfor details. **b-c,** Gold standard FSC curve and euler angle distribution of the HKU5 $S_{DM1}$ protein is estimated by cryoSPARC. **d,** FSC curve of the refined model of HKU5 $S_{DM1}$ protein versus the overall map that it is refined against (black); of the model refined against the first half map versus the same map (red); and of the model refined against the first half map versus the second half map (green). The small difference between the red and green curves indicates that the refinement of the atomic coordinates did not suffer from overfitting. **e,** The cryo-EM map shown in orthogonal views at a contour level of 4 σ of the HKU5 $S_{DM1}$ protein, with protomers displayed in blue, violet, and green, respectively.

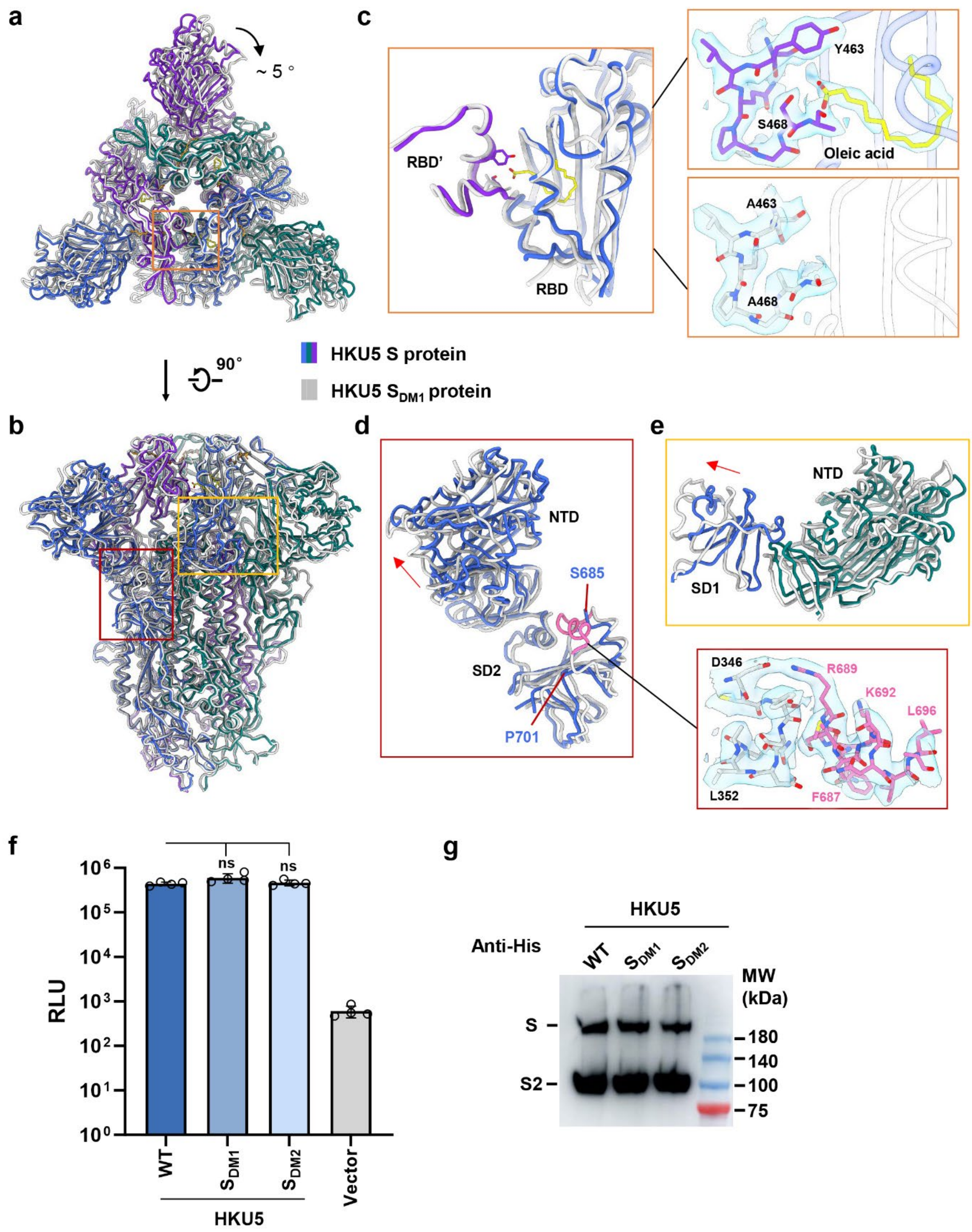


**Supplementary Fig. 5. Structural comparison of the WT HKU5 S protein and the S$_{DM1}$ protein.**

**a-b,** Overall structural comparison between the WT HKU5 S protein and the SDM1 protein, viewed from two orthogonal orientations. **c,** The comparison of the local structure and the cryo-EM map of the WT HKU5 S protein (at a contour level of 6 σ) and the S$_{DM1}$ protein (at a contour level of 6 σ) at the oleic acid binding site. The oleic acid molecule is shown in yellow, and the residues are represented as stick models. The results show that the mutations cause significant changes in the local structure, eliminating the binding of the oleic acid. **d-e,** The local structural changes in the NTD of the HKU5 S protein upon oleic acid binding. Red arrows indicate the direction of structural movement, and pink regions represent the remodeled secondary structures in the HKU5 $S_{DM1}$ protein with the cryo-EM map at a contour level of 6 σ. **f,** Entry of the WT S or the fatty acid binding site mutants ($S_{DM1}$ or $S_{DM2}$) of HKU5 pseudovirus (PSV) into Lenti-X 293T cells stably expressing *Pipistrellus abramus* ACE2. Data represent mean ± SD of four technical replicates from one representative experiment (three independent experiments performed in total). Statistical significance was determined using two-sided unpaired Student's *t*-tests. ns, not significant ($P \geq 0.05$). **g,** Western blot assessment of WT and mutants of HKU5 S cleavage in pseudovirus. Representative images from three independent experiments showing similar results. The uncropped blot is available in Supplementary Fig. 17. Source data are provided as a Source Data file.

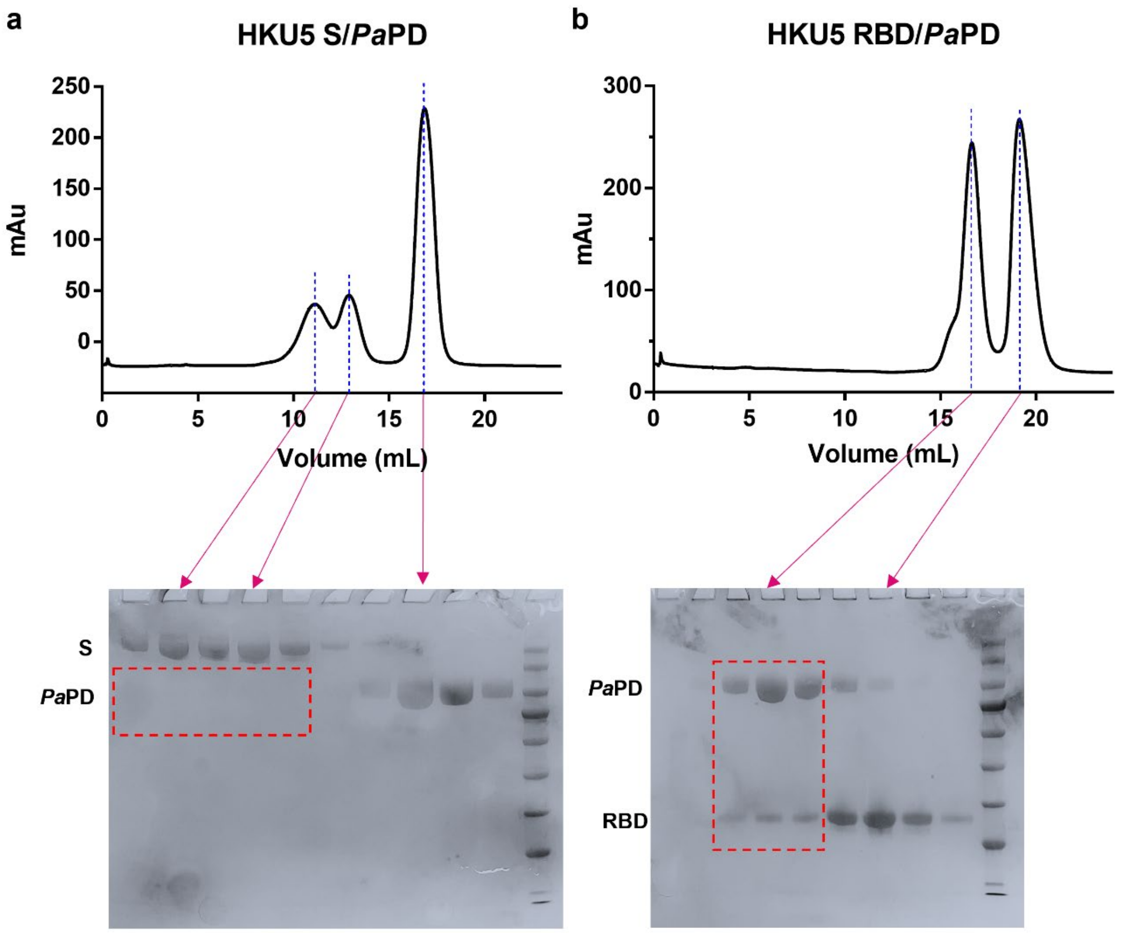


**Supplementary Fig. 6. Gel filtration and SDS-PAGE analysis of the HKU5 S protein or its RBD incubated with *Pa*PD.**

**a,** Top: gel filtration elution profile of HKU5 S protein and *Pa*PD incubated at a molar ratio of 1:5 for 1 hour. Bottom: SDS-PAGE analysis of fractions collected from the gel filtration. The results indicate that HKU5 S protein and *Pa*PD do not co-elute, suggesting that they do not form a stable complex under these conditions. **b,** Top: gel filtration elution profile of HKU5 RBD and *Pa*PD incubated at a molar ratio of 5:1 for 1 h. Bottom: SDS-PAGE analysis of fractions collected from the gel filtration. The results indicate that HKU5 RBD and *Pa*PD can be co-eluted, suggesting that they form a stable complex under these conditions.

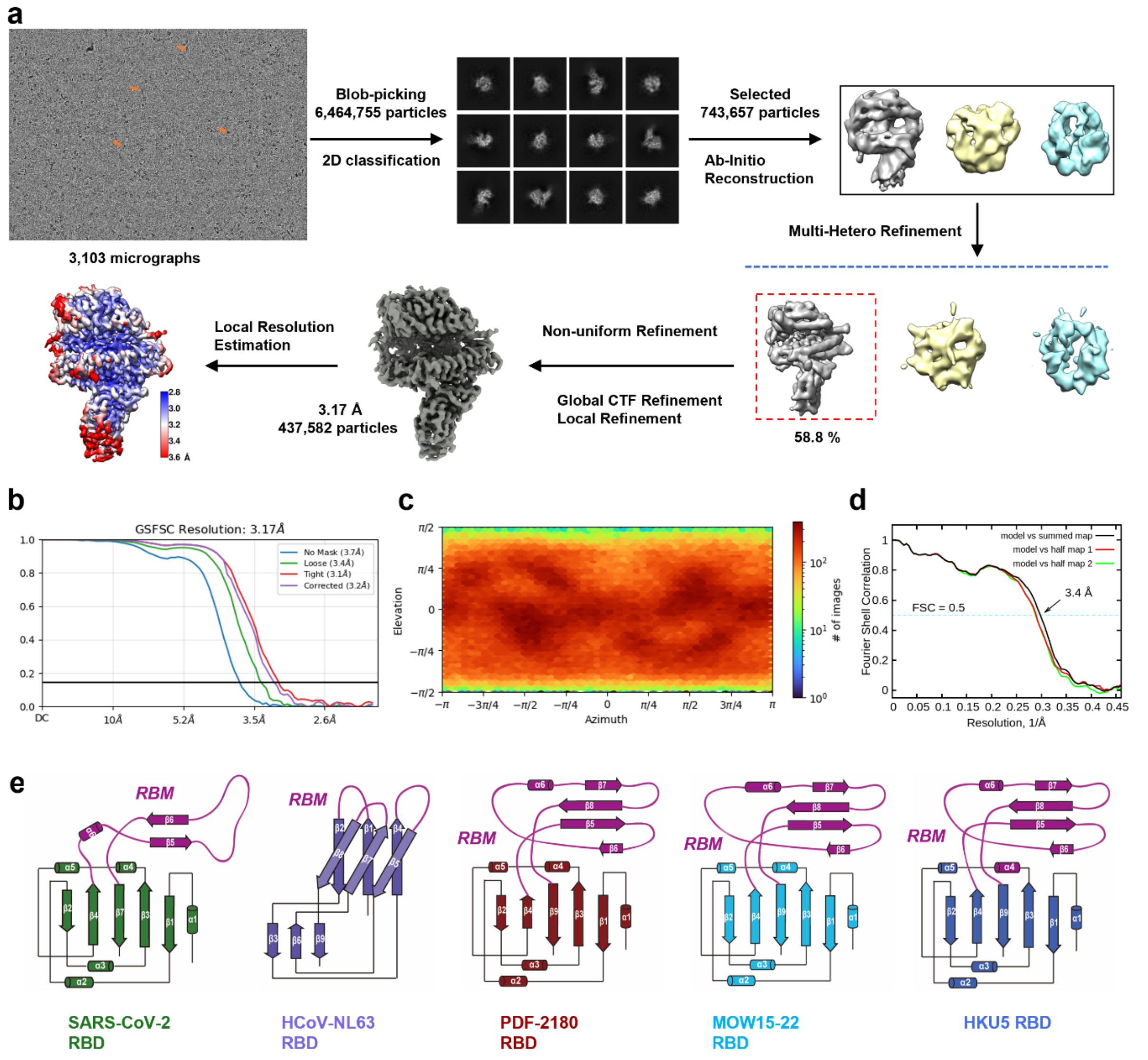


**Supplementary Fig. 7. Cryo-EM data processing and analysis for the HKU5 RBD–*Pa*PD complex.**

**a,** Flowchart of cryo-EM data processing, see the "Data Processing" section in Methods for details. **b-c,** Gold standard FSC curve and euler angle distribution of the HKU5 RBD–*Pa*PD complex is estimated by cryoSPARC. **d,** FSC curve of the refined model of HKU5 RBD–*Pa*PD complex versus the overall map that it is refined against (black); of the model refined against the first half map versus the same map (red); and of the model refined against the first half map versus the second half map (green). The small difference between the red and green curves indicates that the refinement of the atomic coordinates did not suffer from overfitting. **e,** RBD topologies of SARS-CoV-2 (green), HCoV-NL63 (violet), PDF-2180 (red), MOW15-22 (cyan) and HKU5 (marine blue), with receptor-binding motif (RBM) region marked in purple.

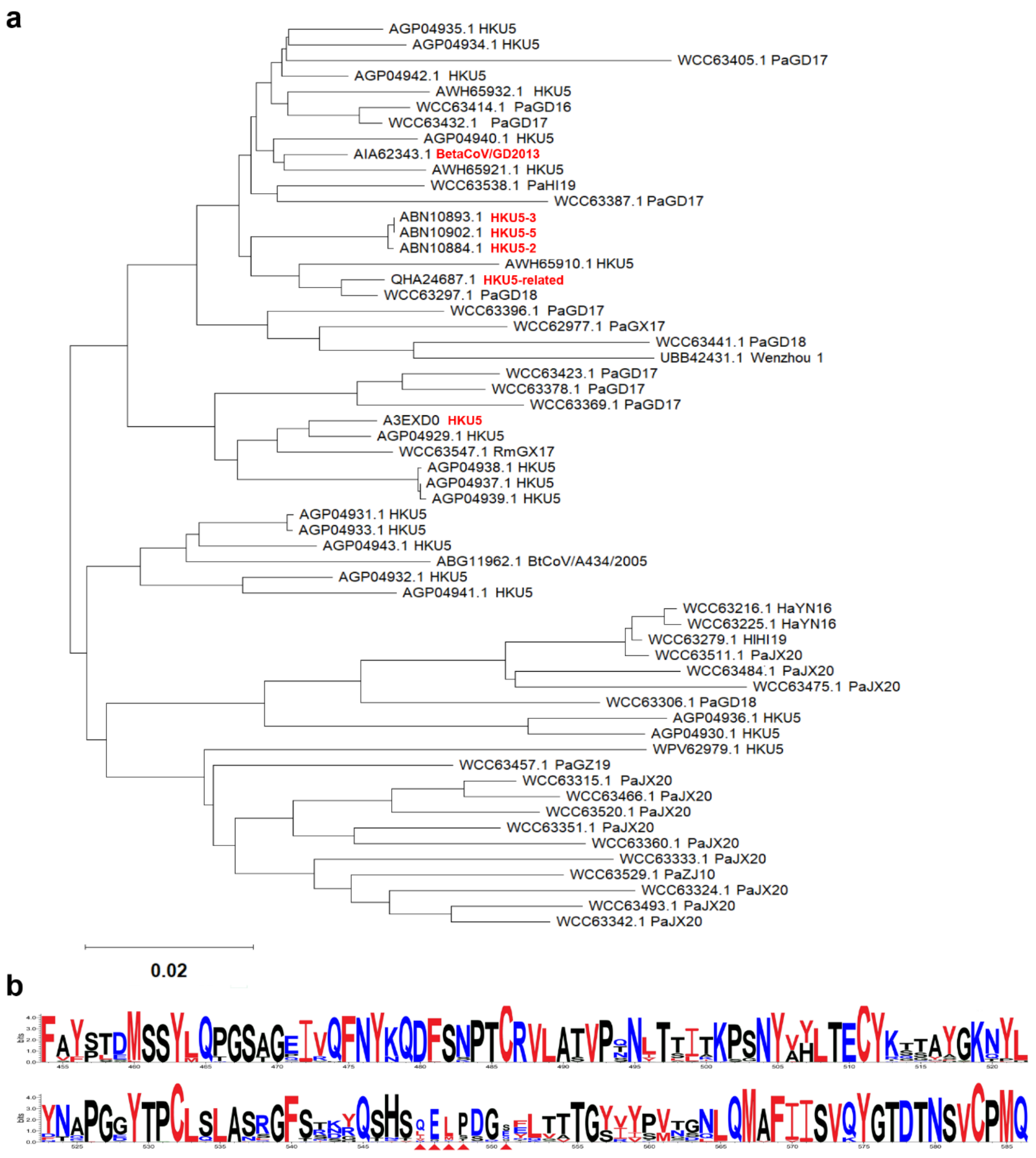


**Supplementary Fig. 8. Phylogenetic analysis and sequence conservation logo of the RBM region of HKU5 lineages.**

**a,** Phylogenetic tree of the S protein sequences in the NCBI protein database with >80% identity to the HKU5 S protein. Phylogenetic tree of HKU5 and related coronaviruses based on RBD sequences was constructed using ClustalW alignment and the Neighbor-Joining method in MEGA 7.0 with 1,000 bootstrap replicates. **b,** The sequence conservation logo with conserved interface residues highlighted. The insertion mutations are marked by solid red triangles. The viruse and protein accession numbers are as follows: Pipistrellus bat coronavirus HKU5 (A3EXD0, AGP04929.1, AGP04937.1, AGP04939.1, AGP04938.1, AGP04942.1, AGP04935.1, AGP04934.1, AGP04940.1, AGP04932.1, AWH65921.1, AGP04931.1, AGP04933.1, AGP04936.1, AGP04943.1, AGP04930.1), BtPa_BetaCoV/GD2013

(AIA62343.1), HKU5-2 (ABN10884.1), HKU5-3 (ABN10893.1), HKU5-related (QHA24687.1), HKU5-5 (ABN10902.1), Bat Coronavirus RmGX17 (WCC63547.1), Bat Coronavirus PaHI19 (WCC63538.1), Bat Coronavirus PaGZ19 (WCC63457.1), Bat Coronavirus PaJX20 (WCC63315.1, WCC63520.1, WCC63529.1), Bat Coronavirus PaGD16 (WCC63315.1, WCC63520.1, WCC63529.1), Bat Coronavirus PaGD17 (WCC63423.1, WCC63378.1, WCC63369.1, WCC63396.1, WCC63414.1, WCC63432.1, WCC63387.1, WCC63351.1, WCC63441.1), and Bat Coronavirus PaGD18 (WCC63297.1, WCC63396.1).

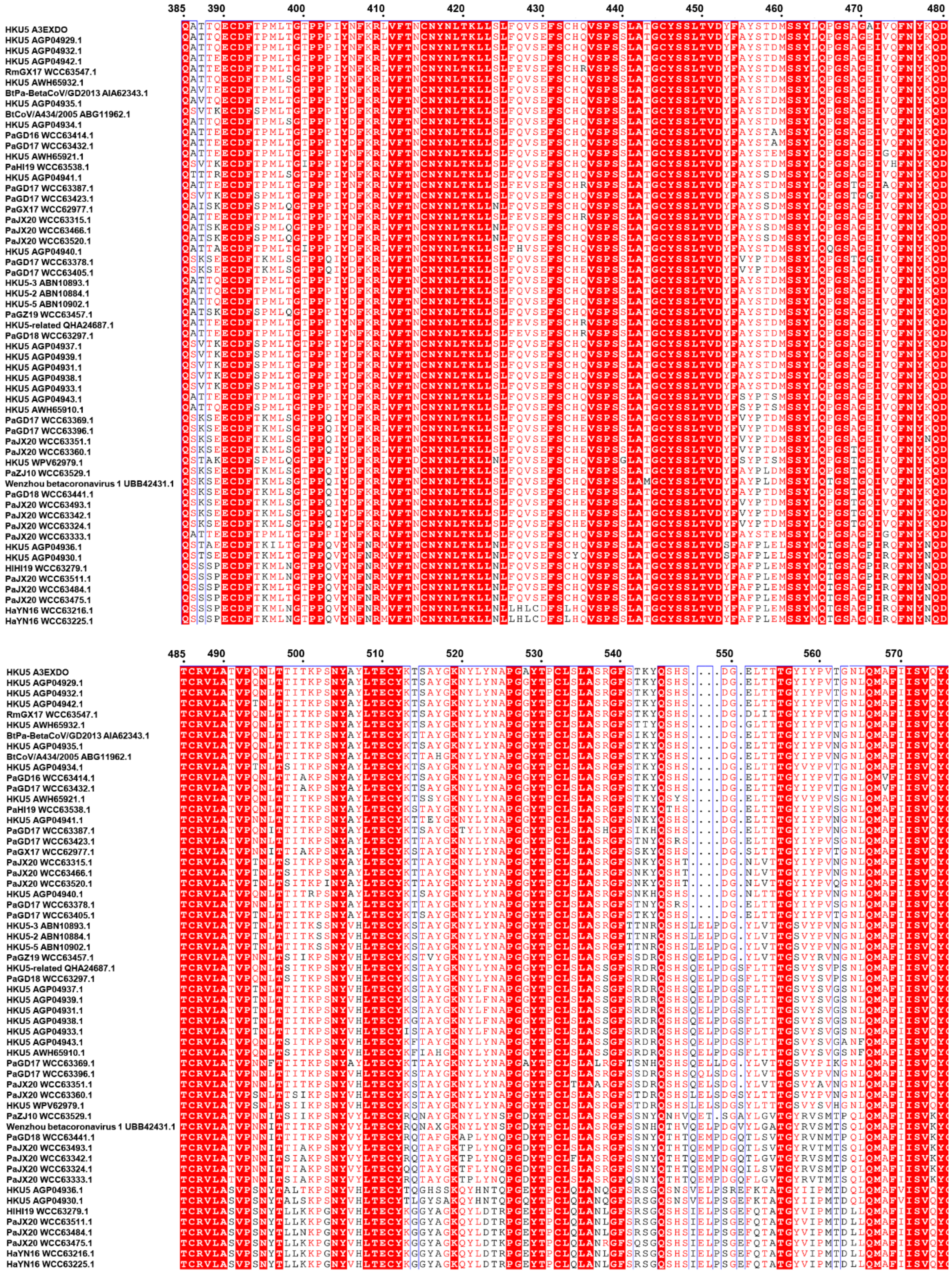

**Supplementary Fig. 9. Multiple sequence alignment of RBD from different HKU5 sub-clades.**

Multiple sequence alignment of the receptor-binding domains (RBDs) from HKU5 and related coronaviruses was performed using Multalin with default settings (https://multalin.toulouse.inra.fr). The specific viruses and their corresponding protein accession numbers are detailed in Supplementary Fig. 8.

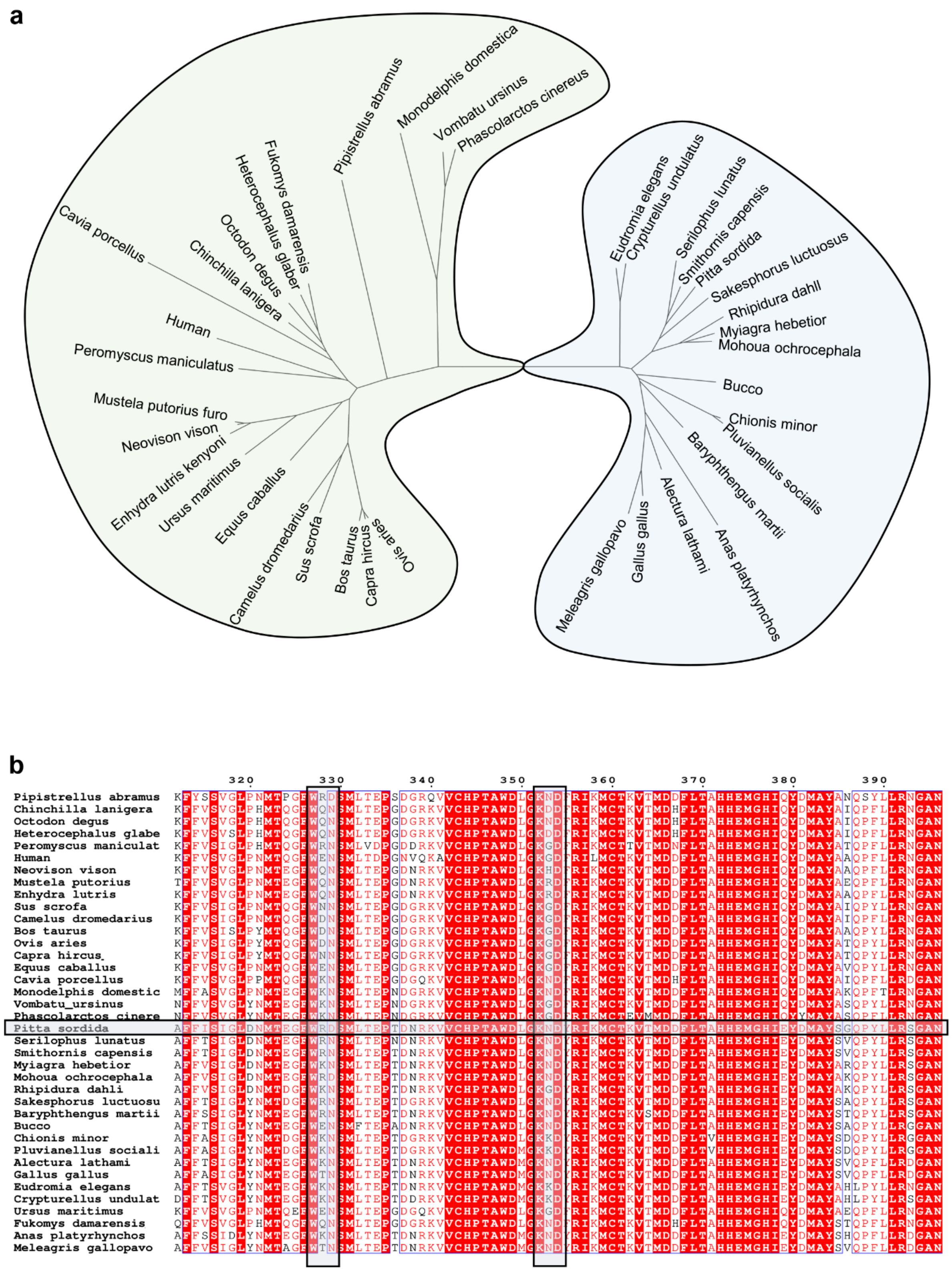


**Supplementary Fig. 10. Evolutionary analysis of *Pipistrellus abramus* ACE2 homologous sequences.**

**a,** Phylogenetic tree of *Pipistrellus abramus* ACE2 homologous sequences. Avian species are marked in light blue, and mammalian species are marked in light green. Amino acid sequences of ACE2 from human, avian species, common farm animals, and potential natural hosts of coronaviruses were retrieved from GenBank. Multiple sequence alignment was performed using ClustalW in MEGA 7.0 with default parameters. A phylogenetic tree was then constructed using the Neighbor-Joining (NJ) method with the Poisson correction model. The reliability of the tree topology was evaluated with 1,000 bootstrap replicates. The final tree was visualized and annotated in MEGA. **b,** Multiple sequence alignment of ACE2 homologous sequences from multiple species was performed using Multalin with default settings. The conserved motifs are indicated by black boxes. The species and protein accession numbers are as follows: *Pipistrellus abramus* (C7ECT9), *Chinchilla lanigera* (A0A8C2UPB0), *Peromyscus maniculatus bairdii* (A0A6I9KY05), *Monodelphis domestica* (F6WXR7), *Pitta sordida* (A0A851FF99), *Myiagra hebetior* (A0A7K9J5U7), *Serilophus lunatus* (A0A7L1DC41), *Baryphthengus martii* (A0A7K9DU94), *Alectura lathami* (A0A7L0W6I4), *Bucco* (A0A7K9IC27), *Eudromia elegans* (A0A7K7VCA6), *Sus scrofa* (K7GLM4), *Bos taurus* (Q58DD0), *Ovis aries* (W5PSB6), *Capra hircus* (A0A452EVJ5), *Equus caballus* (F6V9L3), *Gallus gallus* (A0A5J6CU64), *Anas platyrhynchos* (U3J4G2), *Meleagris gallopavo* (G1NPB8), *Camelus dromedarius* (XP_031301717.1), *Human* (Q9BYF1), *Neovison vison* (A0A8C7BTF2), *Octodon degus* (A0A6P6ESQ8), *Enhydra lutris kenyoni* (A0A2Y9KLV0), *Mustela putorius furo* (Q2WG88), *Heterocephalus glaber* (G5C234), *Vombatu ursinus* (A0A4X2M679), *Phascolarctos cinereus* (A0A6P5M0G0), *Ursus maritimus* (A0A8M1G354), *Fukomys damarensis* (A0A091D044), *Mohoua ochrocephala* (A0A7K7XXX4), *Smithornis capensis* (A0A7K8QUJ4), *Rhipidura dahli* (A0A7K9W0V6), *Sakesphorus luctuosus* (A0A7K8YCL7), *Crypturellus undulatus* (A0A7K4LNH7), *Chionis minor* (A0A7K7FWK2), *Pluvianellus socialis* (A0A7L3D992), *Cavia porcellus* (H0VSF6).

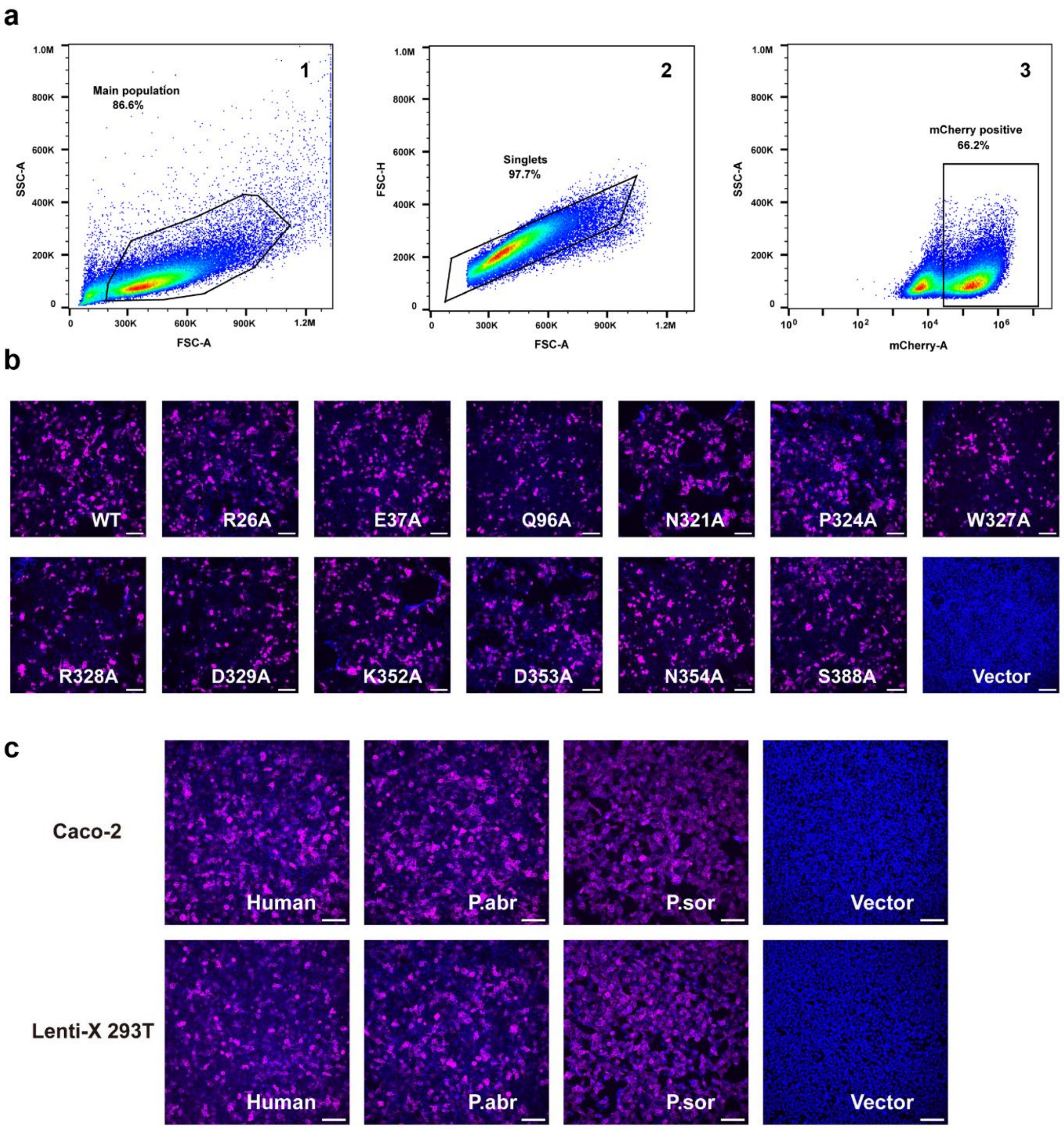


**Supplementary Fig. 11. Gating strategy and expression of ACE2 orthologs and mutants.**

**a,** Flow cytometry-based sorting strategy of cells stably expressing ACE2 orthologs. The first gate was applied to exclude cell debris, followed by the second gate to eliminate adhered or doublet cells. The third gate was used to select mCherry-positive cells. Representative sorting profiles and population percentages are shown for Caco-2 cells stably expressing *Pipistrellus abramus* ACE2. An identical gating strategy was used for all Lenti-X 293T and Caco-2 cell lines stably expressing ACE2 orthologs. **b,** Immunofluorescence analysis of the Lenti-X 293T cells transiently expressing the indicated *Pipistrellus abramus* (P.abr) ACE2 mutants by detecting FLAG tag fused to the constructs. **c,** Immunofluorescence analysis of the Caco-2 and Lenti-X 293T cells stably expressing three ACE2 orthologs, *Pipistrellus abramus* (P.abr), *Pitta* *sordida* (P.sor) and *Homo sapiens* (Human), by detecting FLAG tag fused to the constructs. Scale bars: 100 μm. Representative images from three independent experiments showing similar results.

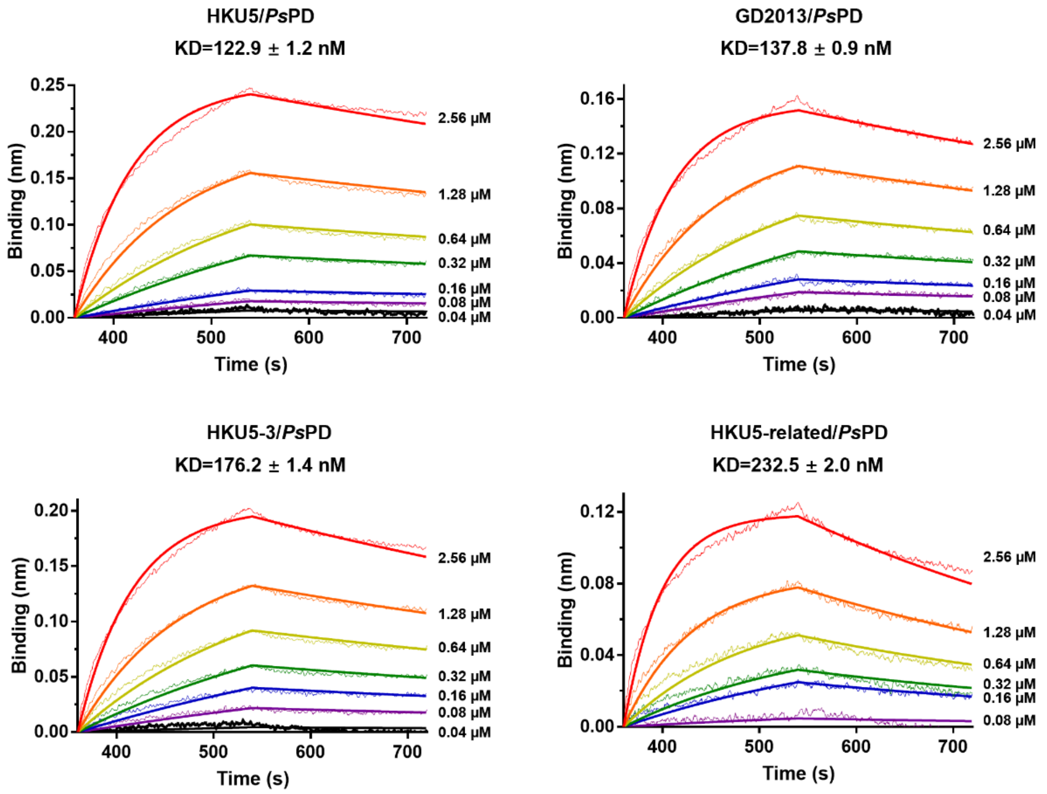


**Supplementary Fig. 12. Interaction analysis of HKU5 sub-clades RBDs with *Ps*PD.**
Binding affinities of RBDs of four HKU5 sub-clades (BatCoV-HKU5 (UniProt ID: A3EXD0), BatCoV-HKU5-3 (GenBank accession: ABN10893.1), BtPa-BetaCoV/GD2013 (GenBank accession: AIA62343.1), and BatCoV-HKU5-Related (GenBank accession: QHA24687.1)) with *Ps*PD, determined using BLI. $R^2$ values are 0.9963, 0.9973, 0.9956, and 0.9892, respectively.

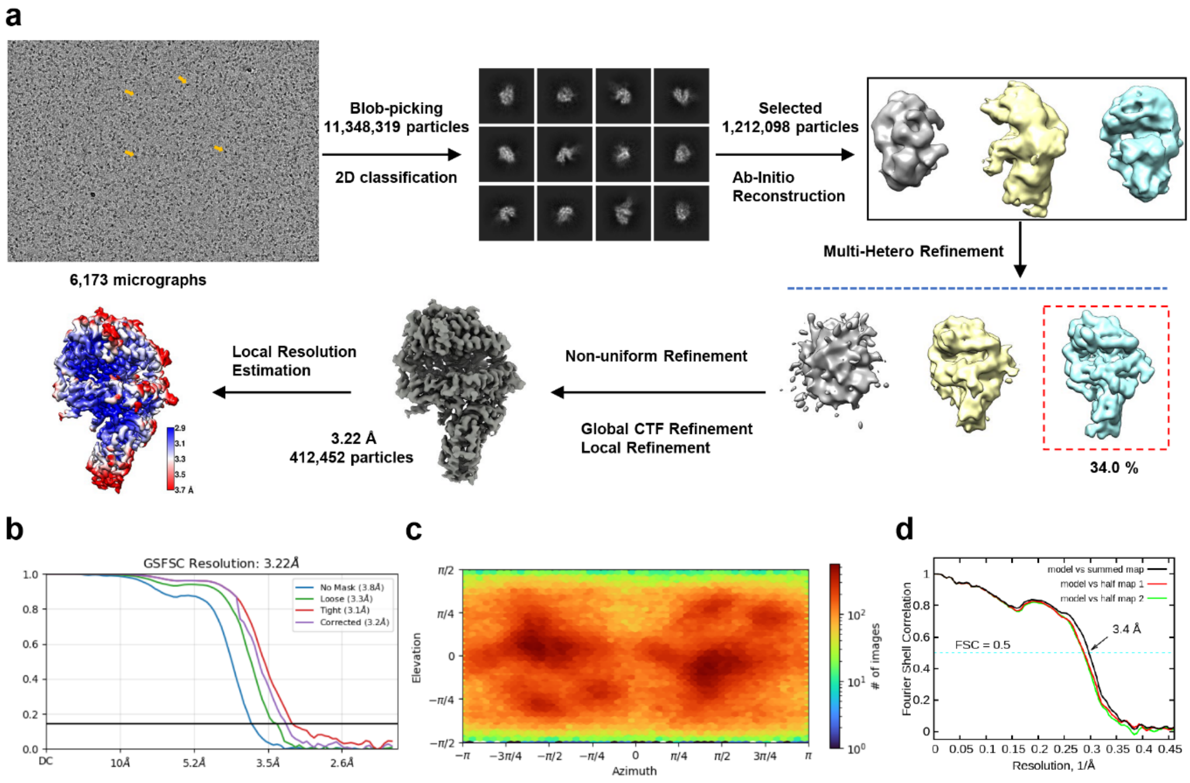


**Supplementary Fig. 13. Cryo-EM data processing and analysis for the HKU5 RBD–*Ps*PD complex.**

**a,** Flowchart of cryo-EM data processing, see the "Data Processing" section in Methods for details. **b-c,** Gold standard FSC curve and euler angle distribution of the HKU5 RBD–*Ps*PD complex is estimated by cryoSPARC. **d,** FSC curve of the refined model of HKU5 RBD–*Ps*PD complex versus the overall map that it is refined against (black); of the model refined against the first half map versus the same map (red); and of the model refined against the first half map versus the second half map (green). The small difference between the red and green curves indicates that the refinement of the atomic coordinates did not suffer from overfitting.

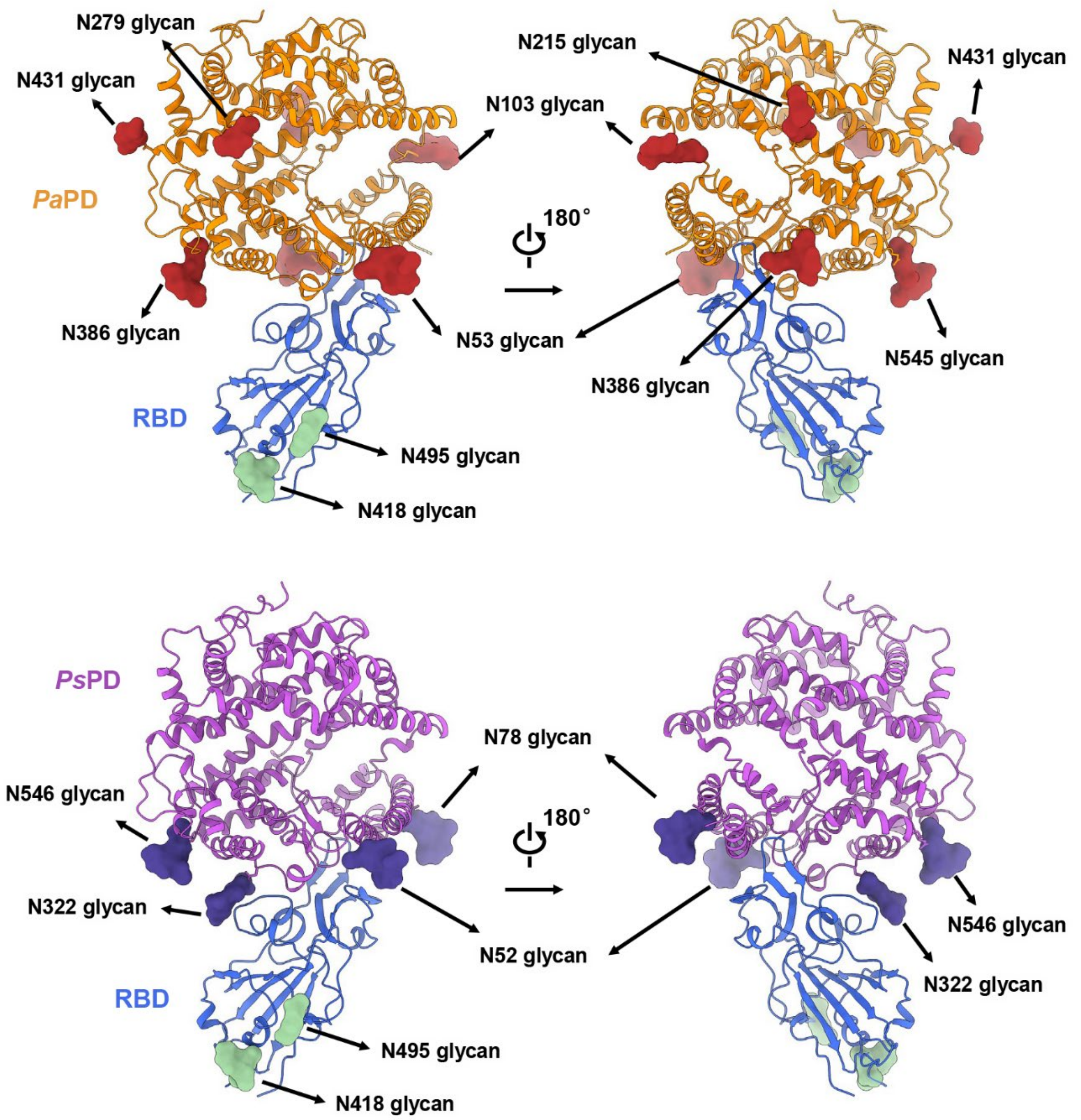


**Supplementary Fig. 14. N-linked glycans of *Pa*PD and *Ps*PD in complex with HKU5 RBD.**

Overall structures of HKU5 RBD (marine blue) bound to *Pa*PD (orange, top panel) and *Ps*PD (purple, bottom panel) in different views, highlighting N-linked glycans displayed as surface representations.

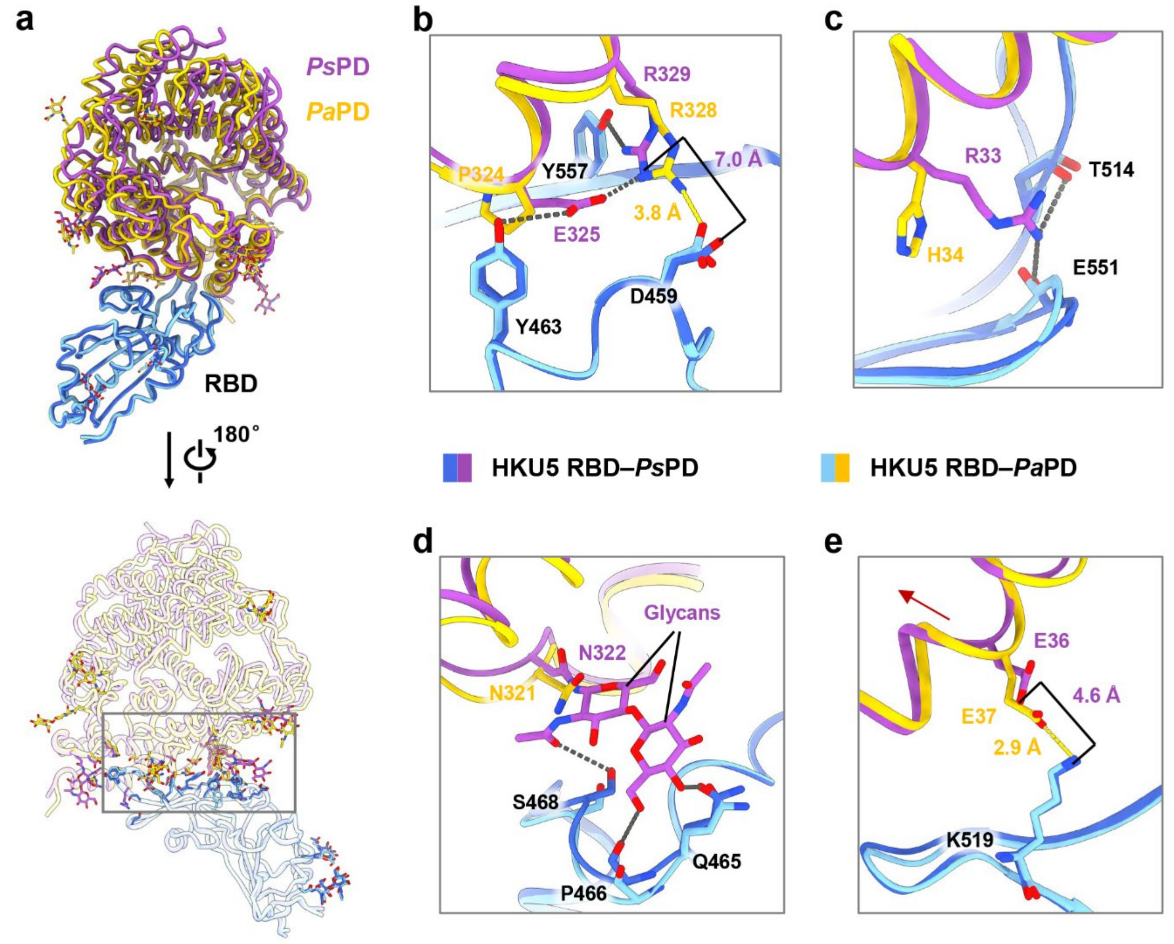


**Supplementary Fig. 15. Comparison of the interfaces between the HKU5 RBD and *Pa*PD or *Ps*PD.**

**a,** Overall structural comparison in different views, with the black box indicating the interaction interface. In the HKU5 RBD–*Ps*PD complex, the RBD and *Ps*PD are shown in marine and purple, respectively. In the HKU5 RBD–*Pa*PD complex, the RBD and *Pa*PD are shown in cyan and gold, respectively. **b-e,** Magnified views of the interface details. Hydrogen bonds are shown as dashed lines, with grey representing RBD–*Ps*PD and yellow representing RBD–*Pa*PD. Disrupted hydrogen bonds in the RBD–*Ps*PD interface, D459–R329 in (**b**) and K519–E36 in (**e**), are annotated with distances.

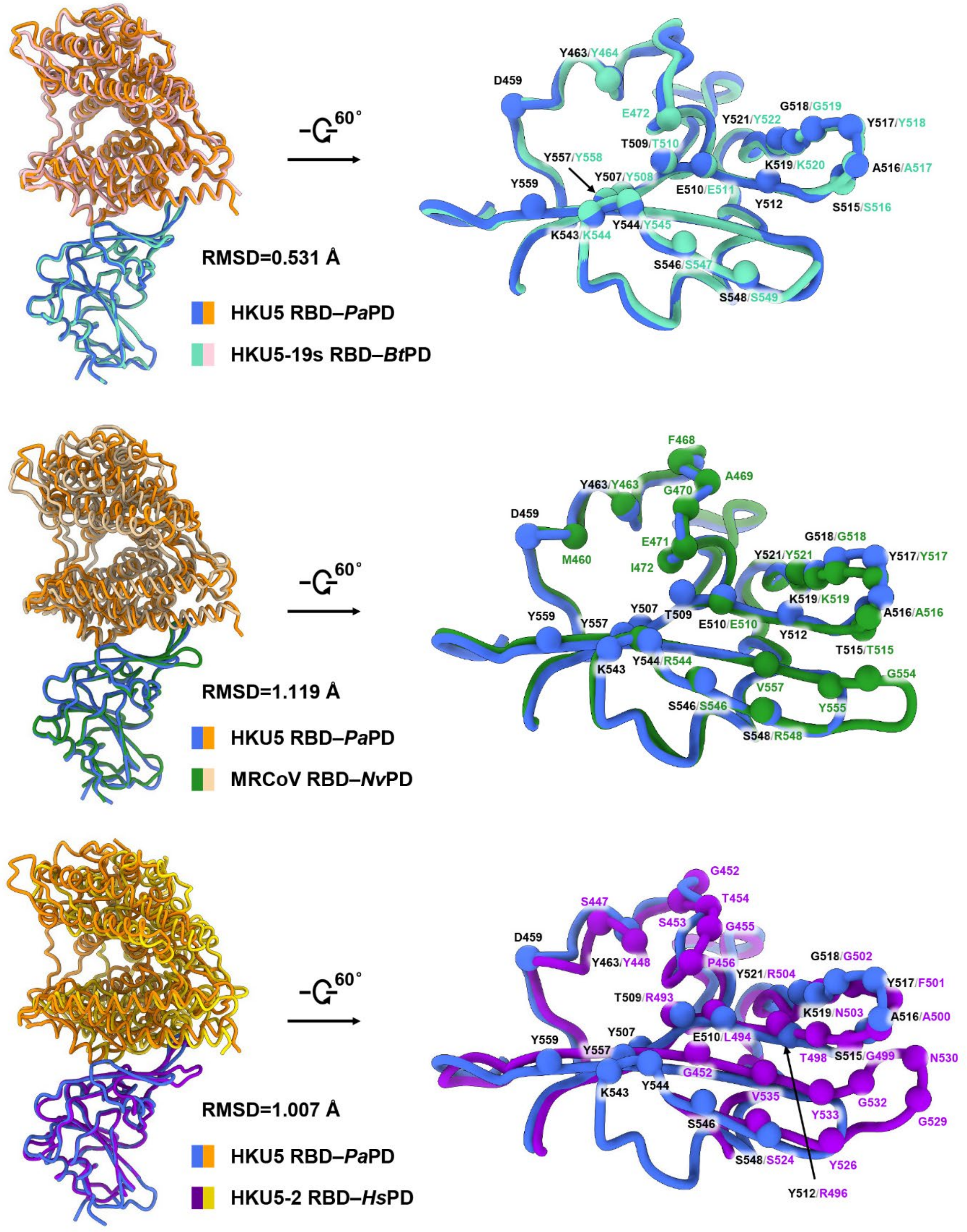


**Supplementary Fig. 16. Structural comparison of available HKU5-related spike proteins.**

Structural comparison of the HKU5 RBD–*Pa*PD complex with HKU5-19s RBD–*Bt*PD (*Bos taurus*; PDB ID: 9E0I), MRCoV RBD–*Nv*PD (*Neogale vison*; PDB ID: 8ZWE), and HKU5-2 RBD–*Hs*PD (*Homo sapiens*; PDB ID: 9JJ6). Residues involved in RBD–ACE2 interactions are shown as spheres in different colors and are labeled accordingly. Residues from HKU5

RBD are labeled in black, from HKU5-19s RBD are in cyan-green, from MRCoV RBD are in forest and those from HKU5-2 RBD are in purple.

**Supplementary Table 1. Characterization of fatty acids associated with the coronavirus S protein.**

| Organism | PDB ID | Ligand ID | Ligand Formula | Ligand MW | Ligand Name | | Ref. |
|---|---|---|---|---|---|---|---|
| HCoV-OC43 | 7SB5 | PLM | C16 H32 O2 | 256.424 | Palmitic acid | 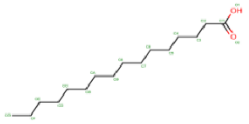 | 35 |
| HCoV-OC43 | 7SBY | 8Z9 | C16 H30 O2 | 254.408 | Sapienic acid | 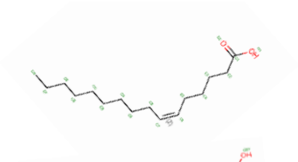 | |
| SARS-CoV-2 | 7E7B | ELA | C18 H34 O2 | 282.461 | 9-Octadecenoic acid | 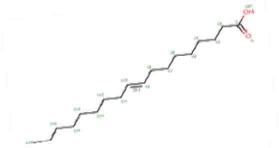 | 52 |
| SARS-CoV-2 | 8H3E | Q83 | C15 H18 N2 O5 | 306.314 | 7-(6-nitro-2,3-dihydroindol-1-yl)-7-oxidanylidene-heptanoic acid | 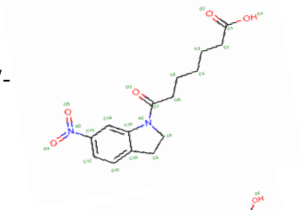 | 37 |
| SARS-CoV-2 | 7Z3Z | STE | C18 H36 O2 | 284.477 | Stearic acid | 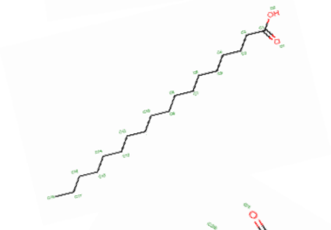 | 53 |
| SARS-CoV-2 | 7Y42 | REA | C20 H28 O2 | 300.435 | Retinoic acid | 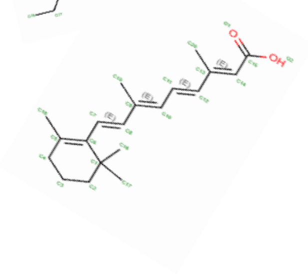 | 54 |
| SARS-CoV | 7ZH1 | EIC | C18 H32 O2 | 280.445 | Linoleic acid | | 55 |
| PCoV_GX | 7CN8 | EIC | C18 H32 O2 | 280.445 | Linoleic acid | 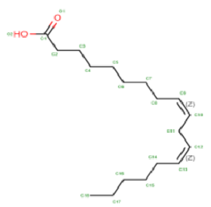 | 56 |
| SARS-CoV-2 | 6ZB4 | EIC | C18 H32 O2 | 280.445 | Linoleic acid | | 17 |
| Bat Coronavirus WIV1 | 8TC0 | EIC | C18 H32 O2 | 280.445 | Linoleic acid | | |
| Civet Coronavirus 007 | 8TC1 | EIC | C18 H32 O2 | 280.445 | Linoleic acid | | 36 |
| Civet Coronavirus SZ3 | 8TC5 | EIC | C18 H32 O2 | 280.445 | Linoleic acid | | |

**Supplementary Table 2. Statistics of cryo-EM data collection, processing, and model refinement.**

| | | | | |
|---|---|---|---|---|
| **Data collection** | | | | |
| EM equipment | Titan Krios (Thermo Fisher Scientific) | | | |
| Voltage (kV) | 300 | | | |
| Detector | Gatan K3 Summit | | | |
| Energy filter | Gatan GIF Quantum, 20 eV slit | | | |
| Pixel size (Å) | 1.087 | | | |
| Electron dose (e-/Å2) | 50 | | | |
| Defocus range (μm) | -1.2 ~ -2.2 | | | |
| Number of collected micrographs | 2,738 | 3,472 | 3,103 | 6,173 |
| Sample | HKU5 S | HKU5 $S_{DM1}$ | RBD-*Pa*PD | RBD-*Ps*PD |
| PDB ID | 9KR8 | 9KR9 | 9KRA | 9KRB |
| EMDB ID | EMD-62522, EMD-64945 (C3 symmetry) | EMD-62523 | EMD-62524 | EMD-62525 |
| **3D Reconstruction** | | | | |
| Software | cryoSPARC | | | |
| Number of used particles | 801,468 | 40,837 | 437,582 | 412,452 |
| Resolution (Å) | 2.39 | 2.80 | 3.17 | 3.22 |
| Symmetry | C1 | | | |
| Map sharpening B factor ($Å^2$) | -91.7 | -88.2 | -190.3 | -187.4 |
| **Refinement** | | | | |
| Software | Phenix | | | |
| Cell dimensions | | | | |
| a=b=c (Å) | 347.84 | 347.84 | 208.704 | 208.704 |
| α=β=γ (˚) | 90 | | | |
| Model composition | | | | |
| Protein residues | 3,489 | 3,552 | 789 | 796 |
| Sugar | 81 | 87 | 15 | 11 |
| Fatty acids | 6 | 3 | 0 | 0 |
| R.m.s deviations | | | | |
| Bonds length (Å) | 0.008 | 0.008 | 0.006 | 0.007 |
| Bonds Angle (˚) | 1.095 | 1.012 | 0.751 | 0.819 |
| Ramachandran plot statistics (%) | | | | |
| Preferred | 93.49 | 93.98 | 93.38 | 91.79 |
| Allowed | 5.96 | 5.93 | 6.11 | 7.95 |
| Outlier | 0.55 | 0.08 | 0.51 | 0.25 |

**Supplementary Fig. 17. Uncropped blot corresponding to Supplementary Fig. 5g.**

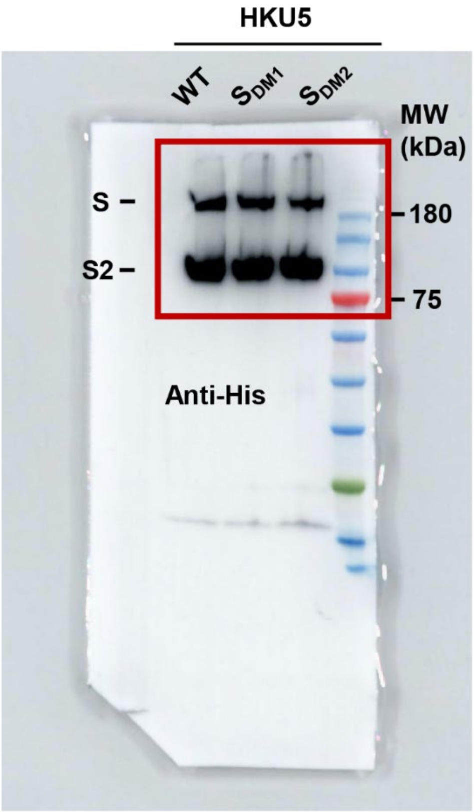
